# DOT1L deletion impairs the development of cortical Parvalbumin-expressing interneurons

**DOI:** 10.1101/2023.01.24.525363

**Authors:** Arquimedes Cheffer, Marta Garcia-Miralles, Esther Maier, Ipek Akol, Henriette Franz, Vandana Shree Vedartham Srinivasan, Tanja Vogel

## Abstract

The cortical plate is composed of excitatory and inhibitory neurons, the latter of which originate in the ganglionic eminences. From their origin in the ventral telencephalon, interneuron precursors migrate during embryonic development over some distance to reach their final destination in the cortical plate. The histone methyltransferase DOT1L is necessary for proper cortical plate development and layer distribution of glutamatergic neurons, however, its specific role on cortical interneuron development has not yet been explored. Here, we demonstrate that DOT1L affects interneuron development in a cell-autonomous manner. Deletion of *Dot1l* in MGE-derived interneuron precursor cells results in an overall reduction and altered distribution of GABAergic interneurons in the cortical plate at postnatal day (P) 0. Furthermore, we observed an altered proportion of GABAergic interneurons in the cortex and striatum at P21 with a significant decrease in Parvalbumin (PVALB)-expressing interneurons. Altogether, our results indicate that reduced numbers of cortical interneurons upon DOT1L deletion results from altered post-mitotic differentiation/maturation.

## Introduction

The development of the mammalian cerebral cortex follows an orchestrated set of events, including cell proliferation, migration and differentiation. Through these events cell types arise in a spatio-temporal dependent manner, including the two main neuronal populations, namely the excitatory glutamatergic neurons and the inhibitory GABAergic interneurons. Glutamatergic neurons originate from the ventricular and subventricular zones of the pallium and migrate along radial glia cells to populate the cortical plate (Rakic 1995). Most of the cortical GABAergic interneurons arise from progenitors localized in the medial and caudal ganglionic eminences (MGE and CGE) in the subpallium (Lavdas et al. 1999; Flames et al. 2007), and a smaller fraction derives from the lateral ganglionic eminence (LGE) (Siddiqi et al. 2021) and the preoptic area (POA) (Gelman et al. 2009). Interneurons migrate tangentially into the cerebral cortex, where they switch to radial migration until they reach their final destination in specific cortical layers (Lavdas et al. 1999; Marín and Rubenstein 2001; Miyoshi and Fishell 2011). Although the cortical GABAergic interneurons, comprising 10-15% of neurons, are much less numerous than their glutamatergic counterparts (85-90%) (Meyer et al., 2011), they are much more diverse regarding morphology, connectivity pattern and electrophysiological properties (reviewed in Tremblay et al. 2016). This makes their integration into neuronal circuitries crucial to ensure a proper cortical excitatory-inhibitory (E-I) balance and a normal brain functioning. Indeed, impairments in GABAergic neurotransmission and consequently altered E-I balance are implicated in many neurological and psychiatric conditions, including Alzheimer’s disease, epilepsy, schizophrenia, autism, depression and mood disorders (reviewed in Nahar et al. 2021; Prévot and Sibille 2021; Ruden et al. 2021). Therefore, information on the cues that govern the appropriate generation, migration, lamination and circuitry integration of GABAergic interneurons gives insight into disease mechanisms, and opens avenues for the development of treatments. In this sense, it has already been shown how transcription factors, cell adhesion molecules, extracellular matrix components, chemoattractive and repulsive factors guide the generation, migration and allocation of GABAergic interneurons in the developing cortex (Liodis et al. 2007; Nóbrega-Pereira et al. 2008; Wester et al. 2019; Limoni et al. 2021).

In addition to the molecules mentioned above, there is an increasing body of evidence showing that epigenetic control of gene expression plays an important role in the development of the central nervous system (CNS) (Juliandi et al. 2010; MuhChyi et al. 2013). Our group has investigated the involvement of the Disruptor of Telomeric Silencing 1-like (DOT1L), a highly conserved histone methyltransferase mediating histone H3 methylation at position lysine 79 (H3K79me), in neuronal differentiation (Büttner et al. 2010; Roidl et al. 2016; Bovio et al. 2019; Franz et al. 2019; Ferrari et al. 2020; Gray de Cristoforis et al. 2020). We demonstrated that loss of DOT1L in forebrain progenitors impairs cortical and hippocampus development. *Dot1l* conditional knockout (*Dot1l* cKO) results in a decreased progenitor pool owing to premature cell cycle exit and premature neuronal differentiation. DOT1L also affects cell fate determination and layer distribution of upper and deep layers (UL and DL) neurons, since *Dot1l-*deficiency leads to decreased numbers of UL neurons and aberrant distribution of DL neurons in the cortical plate (Franz et al. 2019).

The broad expression of DOT1L in the CNS makes it likely that *Dot1l* deletion not only affects the glutamatergic lineage, but also interferes with the development of GABAergic neurons. In fact, we demonstrated that *Dot1l* is necessary for the generation and migration of GABAergic interneurons in the developing spinal cord (Gray de Cristoforis et al. 2020). Furthermore, H3K79me2 marks deposited by DOT1L in the chromatin of GABAergic medium spiny neurons mediate stress response and susceptibility to depression-like behaviors in a mouse model. Notably, intraperitoneal injection of a DOT1L inhibitor reversed the social impairments associated with the depressive phenotype in these mice (Kronman et al. 2021).

Despite emerging evidence, it is not reported as of yet whether and how DOT1L is involved specifically in the development of cortical GABAergic interneurons. Here, we employed bulk RNA sequencing together with histological methods on two different *Dot1l* cKO mice, using either Foxg1^cre^ or Nkx2.1^cre^ driver lines, to reveal DOT1L effects on interneuron development. We show that *Dot1l* deletion impairs the embryonic development of GABAergic interneurons. Specifically, in MGE-derived interneurons we observed sustained effects on the *Pvalb*-expression fraction that exhibited postnatal reduction. Our transcriptomic data suggest altered post-mitotic differentiation/maturation as one possible underlying mechanism.

## Material and Methods

### Animals

Forkhead box G1 (Foxg1)-cre animals (Hébert and McConnell 2000) were mated with floxed *Dot1l*. Animals with the genotype Foxg1^cre/+^;Dot1l^flox/flox^ (cKO) were analyzed in comparison to littermate controls with Foxg1^+/+^;Dot1l^flox/+^ or Foxg1^+/+^;Dot1l^flox/flox^. NK2 Homeobox 1 (Nkx2.1)-cre animals (Xu et al. 2008; JAX #008661) were initially mated with a R26R(EYFP) line (Srinivas et al. 2001; JAX #006148) to generate Nkx2.1-cre/R26R(EYFP) animals, which were subsequently mated with floxed *Dot1l*. Animals with the genotype Nkx2.1^cre/+^/R26R(EYFP);Dot1l^flox/flox^ (cKO) were analyzed in comparison to littermate controls with Nkx2.1^+/+^/R26R(EYFP);Dot1l^flox/+^, Nkx2.1^+/+^/R26R(EYFP);Dot1l^flox/flox^, Nkx2.1^cre/+^/R26R(EYFP);Dot1l^+/+^ or Nkx2.1^cre/+^/R26R(EYFP);Dot1l^flox/+^. Animal experiments were approved by the animal welfare committees of the University of Freiburg and local authorities (G16/11 and G21/0182).

### Mouse brain dissection and fixation

Pregnant females were sacrificed by cervical dislocation and embryos at different embryonic stages (E12.5, E14.5, E16.5 and E18.5) were harvested and brains were dissected in sterile 1X phosphate-buffered saline (PBS). After dissection, brains were fixed in 4% paraformaldehyde (PFA) for 6 to 24 h at 4°C and cryoprotected in 30% sucrose in 1X PBS at 4°C.

### BrdU injections

For BrdU pulse labeling of progenitors, pregnant females were injected at E14.5 with 140 μg/g body weight BrdU (Sigma). Embryos were harvested 1 h later and brains were dissected and fixed in 4% PFA for 6 to 24 h at 4°C followed by cryoprotection with 30% sucrose in 1X PBS at 4°C.

### Intracardiac Perfusion

Mice at 3 weeks of age were anesthetized with intraperitoneal injections of a ketamine (80 mg/kg)/xylazine (10 mg/kg) mixture in isotonic saline solution (0.9% NaCl) with a maximal injection volume of 10 μl/g of body weight. Mice were perfused with ice-cold PBS followed by ice-cold 4% PFA in PBS. Brains were removed and left in 4% PFA for 24 h and then transferred into a 30% sucrose solution in PBS.

### Cryosectioning

PFA fixed brains were embedded in TissueTec (SAKURA) and cryosectioned at 14 μm thickness with a Leica CM3050 S cryostat. Sections were mounted on SuperFrost Plus Microscope slides (Thermo Scientific), immediately dried at 50°C on a heating plate for 1-2 h and then stored at -20°C until needed.

### Immunostaining, imaging and quantification

Slides were dried for 30 min at 37°C, permeabilized with 0.025% Triton-X100/TBS for 45 min and blocked in 1% BSA/TBS for 1 h at room temperature (RT). Primary antibody incubation was done overnight at 4°C in 10% donkey or goat serum and 1% BSA in TBS. Slides were washed 3 times in TBS and incubated with fluorescent secondary antibody in 1% BSA/TBS for 1 h at RT. After washing 3 times with TBS, cells were incubated for 5 min with DAPI (4′,6-diamidino-2-phenylindole, ThermoScientific) and washed 3 more times in PBS. Slides were coverslipped with fluorescent mounting medium (#S3023, DAKO, Jena, Germany). For BrdU immunostaining, slides were incubated with 2N HCl for 30 min at RT, twice with borat buffer (0.1M) for 10 min at RT, and proteinase K digestion in TBS (1:300) for 1 min at RT followed by incubation with 10% normal donkey or goat serum, 1% BSA and 0.3% Triton-X100 in TBS (Blocking solution) for 1 h at RT. First antibody was incubated with blocking solution for 48 h at 4°C. Secondary antibody and DAPI incubation, and coverslip procedures were performed as above. Images were obtained using an Axioplan M2 fluorescent microscope (Zeiss) equipped with an Apotome.2 module or with a Leica Sp8 confocal microscope. Quantification of total number of positive cells in each ROI was done using the cell counter option of Fiji-ImageJ. Total number of cells was normalized to its ROI area (cells/mm^2^). The following first and secondary antibodies were used: anti-cleaved Caspase 3 (rabbit; 1:200; Cell Signaling; 9664S), anti-GFP (chicken; 1:500; Abcam; ab13970), anti-BrdU (sheep; 1/100; Abcam; ab1893), anti-Ki67 (rabbit; 1:250; Abcam; ab15580), anti-SOX6 (rabbit; 1:50; Santa Cruz; sc-20092), anti-rabbit Alexa 594 (donkey; 1:500; Thermo Fisher; R37119), anti-rabbit Alexa 488 (donkey; 1:500; Thermo Fisher; A27034), anti-chicken Alexa 488 (goat; 1:500; Thermo Fisher; A78948), anti-sheep Cy3 (donkey; 1:500; Jackson ImmunoResearch; AB 2315778).

### Single molecule fluorescent *in situ* hybridization (smFISH), imaging, and quantification

smFISH was performed with the RNAscope Multiplex Fluorescent Reagent Kit v2 according to the manufacturer’s instructions (ACD, 323270). Briefly, slides were incubated at 40°C for 45 min in the HybEZ™ oven followed by hydrogen peroxide incubation for 10 min at RT. After washing, slides were boiled for 3 min at 95°C in target retrieval buffer and then washed in distilled water and 100% ethanol. Slides were dried at RT and then, incubated with RNA protease III for 21 min at 40°C. Sections were incubated with pre-warmed smFISH probes diluted in probe diluent at 1:50 for 2 h at 40°C. After washing the slides in the wash buffer (WB), slides were incubated with AMP1 and AMP2 for 30 min each at 40°C followed by AMP3 incubation for 15 min at 40°C. The following steps were repeated for each channel used in the probes: HRP was added for 15 min at 40°C followed by incubation with fluorophore diluted in TSA for 30 min at 40°C and HRP blocker for 15 min at 40°C. Finally, sections were counterstained with DAPI (1:1000 in 1X PBS) for 5 min and coverslipped with fluorescent mounting medium (#S3023, DAKO, Jena, Germany). Images were obtained using an Axioplan M2 fluorescent microscope (Zeiss) equipped with an Apotome.2 module or with a Leica Sp8 confocal microscope. Fluorescent RNA molecules were quantified using a set of Fiji-ImageJ scripts described previously (Wang el at., 2019). Briefly, the script “estimatedotradius.ijm” was first run to determine the average dot radius followed by the script “thersholdoptimization.ijm” to determine manually the threshold of each region of interest (ROI) to be processed. The number of RNA molecules in each ROI was normalized to its area (molecules/mm^2^). For animals at P21, quantification of *Pvalb, Npy* and *Sst* RNA molecules in each ROI was done using the cell counter option of Fiji-ImageJ since at this maturation stage RNA molecules were accumulated within cells. Total number of cells were normalized to their corresponding ROI area (cells/mm^2^). The following smFISH probes and fluorophores were used: *Plcdx3*-C3 probe (mouse; ACD; 498091) with Opal 570 fluorophore (1:1500), *Phlda1-*C2 probe (mouse; ACD; 593561) with Opal 570 fluorophore (1:1500), *Pvalb*-C1 probe (mouse; ACD; 421931) with Opal 520 fluorophore (1:800), *Npy-*C2 probe (mouse; ACD; 313321) with Opal 650 fluorophore (1:800) and *Plcdx3*-C3 probe (mouse; ACD; 404631) with Opal 570 fluorophore (1:1500).

### *In situ* hybridization, imaging, and quantification

Probes for in situ hybridization were made by cloning PCR products into pGemTeasy (Promega). 1 μg of the linearized plasmid was transcribed *in vitro* using NTP labelling mix and T7 or SP6 RNA Polymerase, followed by purification with mini-Quick spin RNA columns (Roche). Probes were diluted in hybridization buffer in 1:100 or 1:500 ratio and incubated on the sections overnight at 70°C. After washing and blocking in lamb serum in MABT buffer (5X MAB, 0.1% Tween 20), sections were incubated with Anti-DIG-AP (Roche) at 4°C overnight. After washing, sections were developed with NBT/BCIP (Roche) overnight and mounted. Bright field images were obtained using an Axioplan M2 microscope (Zeiss). Quantification of total number of cells expressing each ISH probe in each ROI was done using the cell counter option of Fiji-ImageJ. Total number of cells was normalized to their corresponding ROI area (cells/mm^2^). The ISH probe sequences were obtained from the following sources: *Nkx2*.*1* (Addgene; 15540), *Lhx6* (Allen Brain Atlas; Experiment #100047358), *Nr2f2* (Allen Brain Atlas; Experiment #100057642), *Prox1* (Addgene; 87129), *Sp8* (Kawakami et al. 2004) and *Sst* (Asgarian et al. 2019).

### RNA isolation from tissue

Pregnant females were sacrificed by cervical dislocation and embryos at E14.5 were harvested, and ventral and dorsal telencephalon were separated. Samples were frozen in liquid nitrogen immediately after dissection and stored at -80°C until RNA extraction. Total RNA was isolated using the RNeasy RNA isolation kit (Qiagen) according to the manufacturer’s instructions. An on-column DNAse digestion was routinely performed. Isolated RNA was kept at -80°C until following qRTPCR experiments.

### Reverse transcription and qRTPCR

1 μg of total RNA was reverse transcribed using RevertAid MMuLV reverse transcriptase kit (Fermentas, Thermo Scientific). qRTPCR analysis was performed on a CFX-Connect Real-Time PCR detection system (Bio-Rad) using GoTaq qPCR Master Mix (Promega). Primers used had an efficiency level between 85% and 110%. qRTPCR results were analyzed using the ΔΔCt method with GAPDH as internal standard. The primers for the qRTPCR target regions used are listed in Supplementary Table 1.

### RNA Sequencing

Total RNA extracted from mouse ventral and dorsal telencephalon was used for RNA sequencing (RNA-seq). Libraries were prepared using the NEBNext Ultra RNA Library Prep Kit for Illumina, following the manufacturer’s instructions. The procedure included depletion of rRNA prior double-stranded cDNA synthesis and library preparation. Samples were sequenced on Illumina HiSeq3000 as paired-end 75 bp reads.

### ChIP sequencing of H3K79 dimethylation

To define the H3K79 dimethylation (H2K79me2) profiles of interneuron gene markers, we carried out a survey on previously published ChIP-seq data (Franz et al. 2019) and analyzed them as described in Ferrari et al. 2020. Briefly, the normalized RPKM means of individual replicates for the cortex of control mice at E14.5 were calculated with deepTools2 bigwigCompare, and the DeepTools2 pyGenomeTracks v3.5 was used to plot the reads over the displayed genomic regions (Ferrari et al. 2020).

### Bioinformatics analysis of RNA-sequencing

Quality Control, trimming and mapping of the RNA-seq fastq files was done on the Galaxy platform using FastQC (Galaxy version 0.73), Cutadapt (Galaxy version 4), and RNA STAR (Galaxy Version 2.7.8a), respectively (Afgan et al. 2022). Adapter trimming and quality filtering was done using Cutadapt with the following configurations: -j 8 -e 0.1 -q 16 -O 3 --trim-n --minimum-length 25 -a AGATCGGAAGAGC. Trimmed and quality filtered RNA-Seq reads were aligned to mm10 genome assembly in RNA STAR with the default configurations for paired-end reads.

Gene level counts were obtained by running featurecounts (Galaxy Version 2.0.1+galaxy2) with default configurations. Differential expression analysis was done using DESeq2 (1.34.0) on R (4.2.0) on count matrix output from featurecounts (Love et al. 2014). Data from wildtype (WT) replicate 4 was excluded from further analysis due to high variability. A linear model controlling for batch effects (∼sex+ litter + genotype) was used and normal log2(Fold Change) shrinkage was applied. Gene counts were VST normalized. GO term enrichment analyses were done using clusterProfiler (4.2.2) (Wu et al. 2021). Visualizations of volcano plots and heatmaps were done using EnhancedVolcano (1.12.0) and pheatmap (1.0.12) packages, respectively (Kolde 2019; Blighe K, Rana S 2022).

### Statistical analyses

For statistical analysis an unpaired, two tailed Student’s *t*-test with Welch’s correction was performed with GraphPad Prism software (version 9.1.1).

### Availability of data and materials

The raw sequencing files were deposited to the NCBI Gene Expression Omnibus (GEO) with accession number GSE221994. All other data types and codes recreating the analyses from the data files can be found at https://github.com/Vogel-lab/DOT1L-interneuron-development as R markdown files, and Galaxy workflows.

## Results

### *Dot1l* deletion in the Foxg1-cre lineage impairs cortical interneuron development

*Foxg1* is involved in interneuron development and we used the recently described Foxg1^cre^ Dot1l cKO mouse line to study the implications of DOT1L in the development of GABAergic interneurons (Franz et al. 2019). Transcriptomic analysis of the dorsal telencephalon of cKO (Foxg1^cre/+^Dotl1^flox/flox^) and control (Foxg1^+/+^Dotl1^flox/+^) mice at E14.5 revealed altered expression of 2106 genes in total (Franz et al. 2019). We manually curated current literature on interneuron development to short list genes expressed by interneurons (Wonders and Anderson 2006; Faux et al. 2012; Chen et al. 2017; Mayer et al. 2018; Mi et al. 2018; Asgarian et al. 2022), which summed up to 534 genes (Supplementary table 2). 86 of these genes expressed by interneurons were differentially expressed upon *Dot1l* cKO (*p. adj. value* < 0.05), with both increased and decreased expression levels when compared to control mice (**Fig. 1A-C**). Notably, the GABAergic markers *Lhx6, Sox6, Sst* and *Npy* decreased significantly upon *Dot1l* deletion, suggesting that the MGE-derived interneuron lineages were affected (**Fig. 1C**). Functional GO-term analysis indicated that genes impacting the maturation, migration and terminal differentiation of interneurons were altered upon DOT1L deletion (**Fig. 1D**).

**Figure 1.**
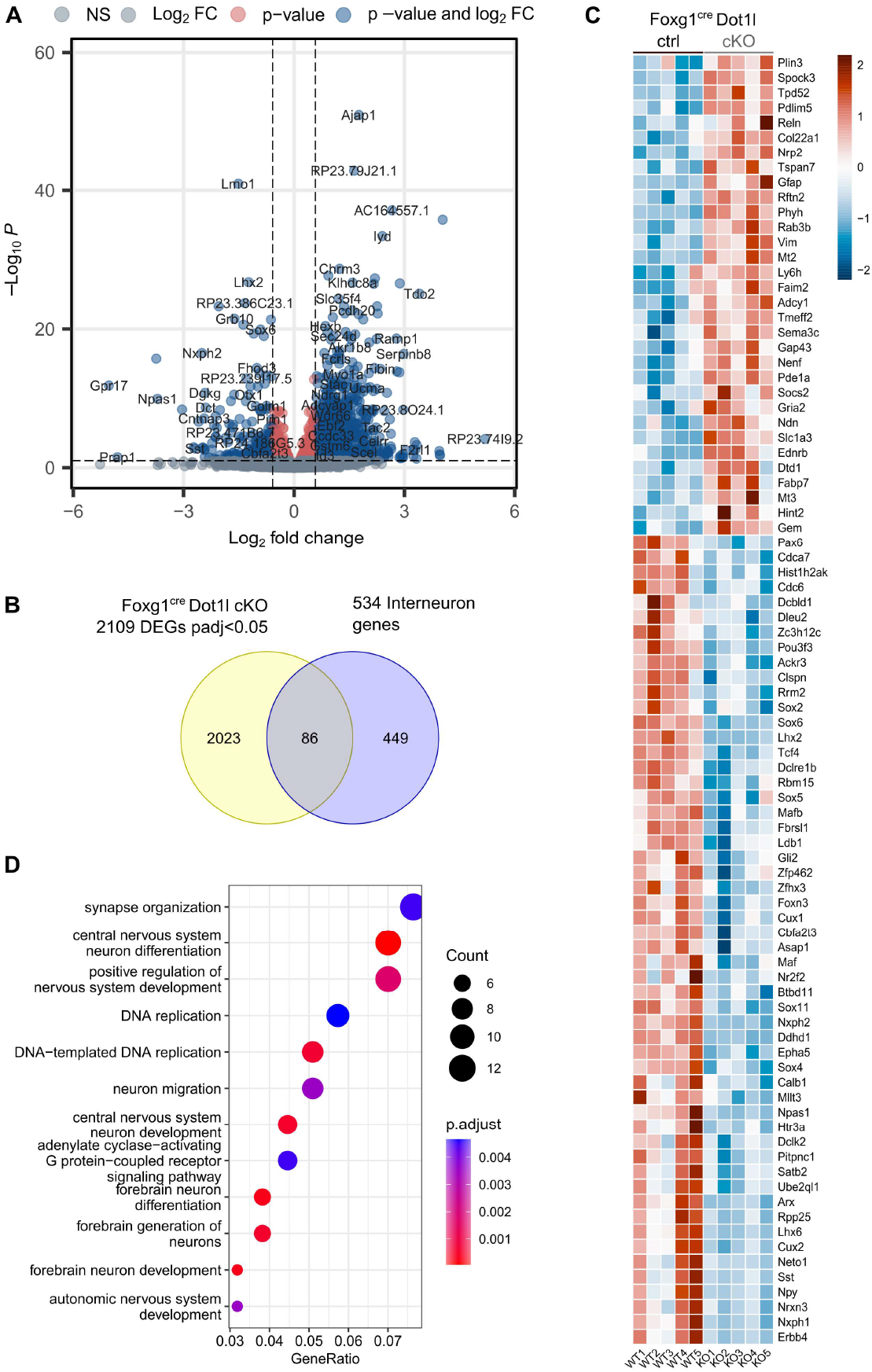
*Dot1l* deletion in the Foxg1-cre lineage impairs cortical interneuron development. **(A)** Volcano plot displaying differentially expressed genes (DEGs) in the dorsal telencephalon of Foxg1^cre^ Dot1l cKO animals compared to controls at E14.5. The y-axis corresponds to the adjusted p-value and the x-axis displays the log_2_Fold Change (log_2_FC). Grey: Transcripts with insignificant adjusted p-values (p > 0.05). Red: Transcripts with differential expression of less than ± log_2_ (0.5). Blue: Transcripts with differential expression of more than ± log_2_ (0.5). Positive log_2_FC represents increase; negative log_2_FC represents decrease in expression upon *Dot1l* deletion. **(B)** Venn diagram of DEGs in dorsal telencephalon of Foxg1^cre^ Dot1l cKO animals at E14.5 that are known to be interneuron gene markers. **(C)** Heatmap of 86 genes at the intersection in **(B)**. DEGs with increased and decreased expression are represented in red or blue, respectively. Scale shows scaled VST normalized gene counts. Data represents sex- and batch-corrected gene expression in forebrain of Foxg1^cre^ Dot1l control (n=5) and cKO (n=5) animals. **(D)** Dotplot of GO term enrichment analysis of DEGs in Foxg1^cre^ Dot1l cKO mice depicting the top 12 significantly enriched biological processes. Total number of genes per group and adjusted p-values are indicated on the right and gene ratios are on the x-axis. Blue depicts the least significantly enriched pathway, and red the most significantly enriched pathway.

*Foxg1* is expressed in the dorsal and ventral telencephalon and resulted in loss of *Dot1l* transcription in both regions (**Fig. 2A**). Taking this broad expression and presence of interneurons in both regions into account, we validated expression of selected genes that impact interneurons separately in the two regions at E14.5 using qRTPCR (**Fig. 2B-H**). Based on the developmental stage in which the interneuron marker genes are expressed (Wonders and Anderson 2006), we categorized the markers according to cell cycle exit, interneuron specification, immature, migrating and mature interneurons. Genes that did not fall into these categories were considered as other interneuron markers. In line with the transcriptomic analysis (**Fig. 1B**), we observed an altered expression of several interneuron markers in both dorsal and ventral telencephalon. Notably, interneuron markers transcribed in immature interneurons, namely *Lhx6, Cux2, Lmo1* and *Sox6*, decreased in both regions (**Fig. 2E**), whereas transcriptional decrease of markers in migrating and mature interneurons only significantly decreased in the dorsal telencephalon, namely *Nrp1, Hrt3a, Erbb4, Sst, Nxph1* and *Npy* (**Fig. 2F, G**). The altered expression levels of these marker genes indicated that (i) Foxg1^cre^ Dot1l cKO affected the MGE-derived Nkx2.1 SST/NPY/PVALB lineage, and (ii) the generation and early specification of interneurons was probably less affected than subsequent maturing stages of development. Confirming the hypothesis of a compromised development of the MGE-derived interneurons was the decreased expression of *Nkx2*.*1*, in contrast to the *Dlx* family members (**Fig. 2D**). Whereas decreased expression of genes characterizing the MGE-derived lineage was prominent, we observed increased expression of CGE-lineage marker genes *Calb2, Ache* and *Reln* in the dorsal telencephalon (**Fig. 2G**).

**Figure 2.**
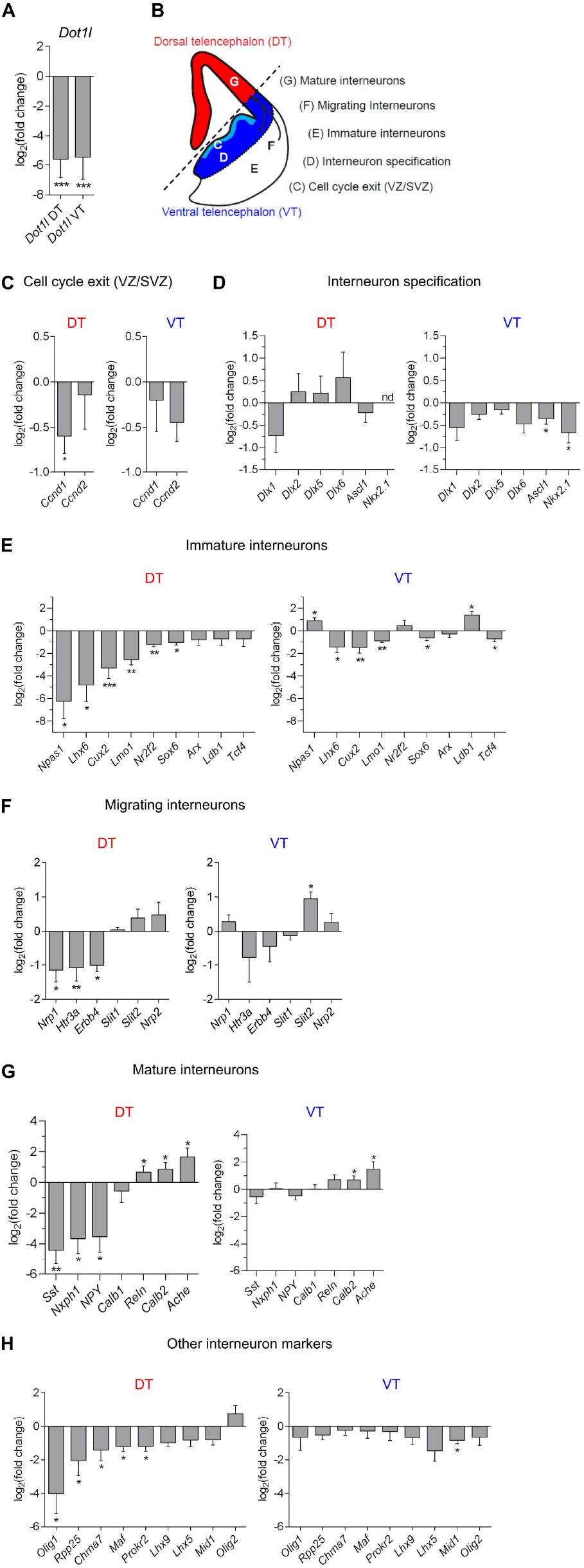
Foxg1^cre^ Dot1l cKO alters expression of interneuron markers in the dorsal and ventral telencephalon. **(A)** qRTPCR of *Dot1l* expression in cKO versus control in dorsal (red, DT) and ventral (blue, VT) telencephalon. Data is represented as the mean ± SEM; n=4-5 (Foxg1^cre^ Dot1l ctrl and cKO animals); \*\*\**P* < 0.001 by unpaired, two tailed Student’s *t*-test with Welch’s correction. **(B)** Schematic view of the FOXG1 expression domains in dorsal (red) and ventral (blue and light blue) telencephalon, and indication of specific regions associated with different stages of interneuron development from C-G. **(C-H)** qRTPCR results for genes falling in the categories C-G, as indicated in (B), and unclassified interneuron markers in (H). Data is represented as the mean ± SEM; n=4-5 (Foxg1^cre^ Dot1l ctrl and cKO animals); \**P* < 0.05, \*\**P* < 0.01, and \*\*\**P* < 0.001 by unpaired, two tailed Student’s *t*-test with Welch’s correction. nd - not detected.

To gain insight into one possible molecular mechanism by which DOT1L regulates interneuron development, we inspected enrichment of H3K79me2 marks at the different interneuron marker genes from a data set derived from the E14.5 wildtype dorsal telencephalon (**Fig. 3A-F)** (Franz et al. 2019). In this set of candidate genes we identified several genes that were marked by H3K79me2 and which therefore could be directly controlled by DOT1L’s enzymatic activity. We found strongest enrichment of genes carrying H3K79me2 in immature interneuron marker genes (**Fig. 3C**). Notably, this state during interneuron development also showed decreased expression of the respective markers in the transcriptome. Using *in situ* hybridization (ISH) or immunohistochemistry (IHC) we observed *in vivo* the reduced expression of *Nkx2*.*1, Lhx6, Nr2f2, Sp8, Sst* and SOX6 (**Fig. 4A-H**), and reduced cell numbers for *Sst* (**Fig. 4C, G**) in the cerebral cortex of *Dot1l* cKO mice at E18.5 compared to the control group. ISH of the MGE markers *Nkx2*.*1* and *Nkx6*.*2*, and of the CGE markers *Prox1* and *Sp8* at the earlier developmental stages E12.5 and E14.5 retrieved positive signals in the expected location (**Fig. S1**). This finding indicated that the alterations observed at E18.5 were not due to gross anatomical misspecification of the MGE and CGE, but more likely due to maturation defects upon DOT1L deletion. Taken together, the data suggested that proper expression of *Dot1l* is necessary for correct development of MGE- and CGE-derived interneurons.

**Figure 3.**
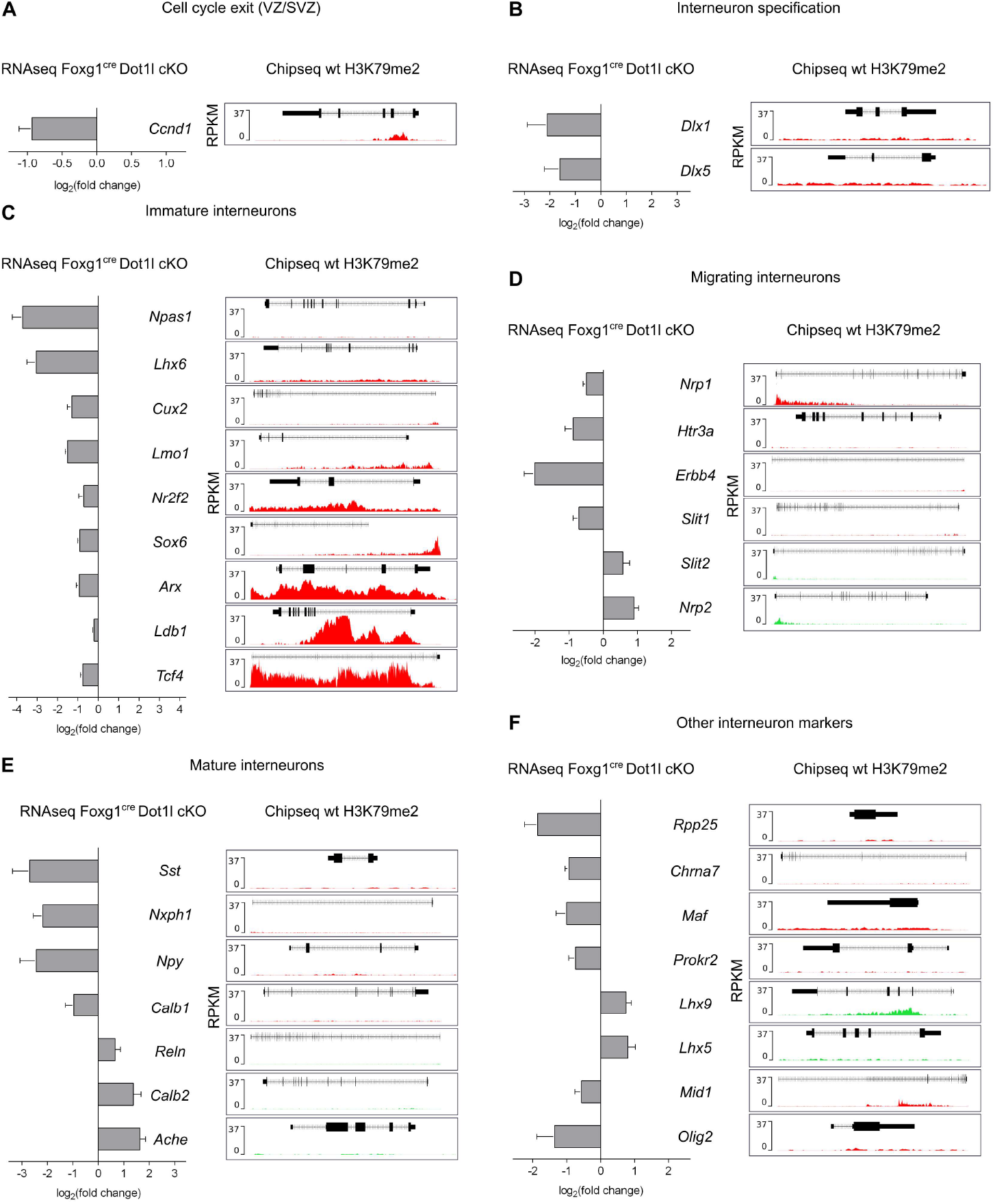
Correlation of transcriptional changes in RNA-seq and H3K79me2 ChIP-seq profile for interneuron markers from E14.5 dorsal telencephalon upon *Dot1l* deletion in the Foxg1-cre lineage. **(A-F)** Left: RNA-seq results (log_2_Fold Change) of Foxg1^cre^ Dot1l cKO animals. Right: ChIP-seq profiles of the same gene shown on the left from the E14.5 dorsal telencephalon of WT animals. Red and green peaks indicate enrichment for H3K79me2 for genes with increased (green) or decreased (red) expression, respectively. Genes are grouped in categories of different developmental stages of interneurons as classified in Fig. 2B.

**Figure 4.**
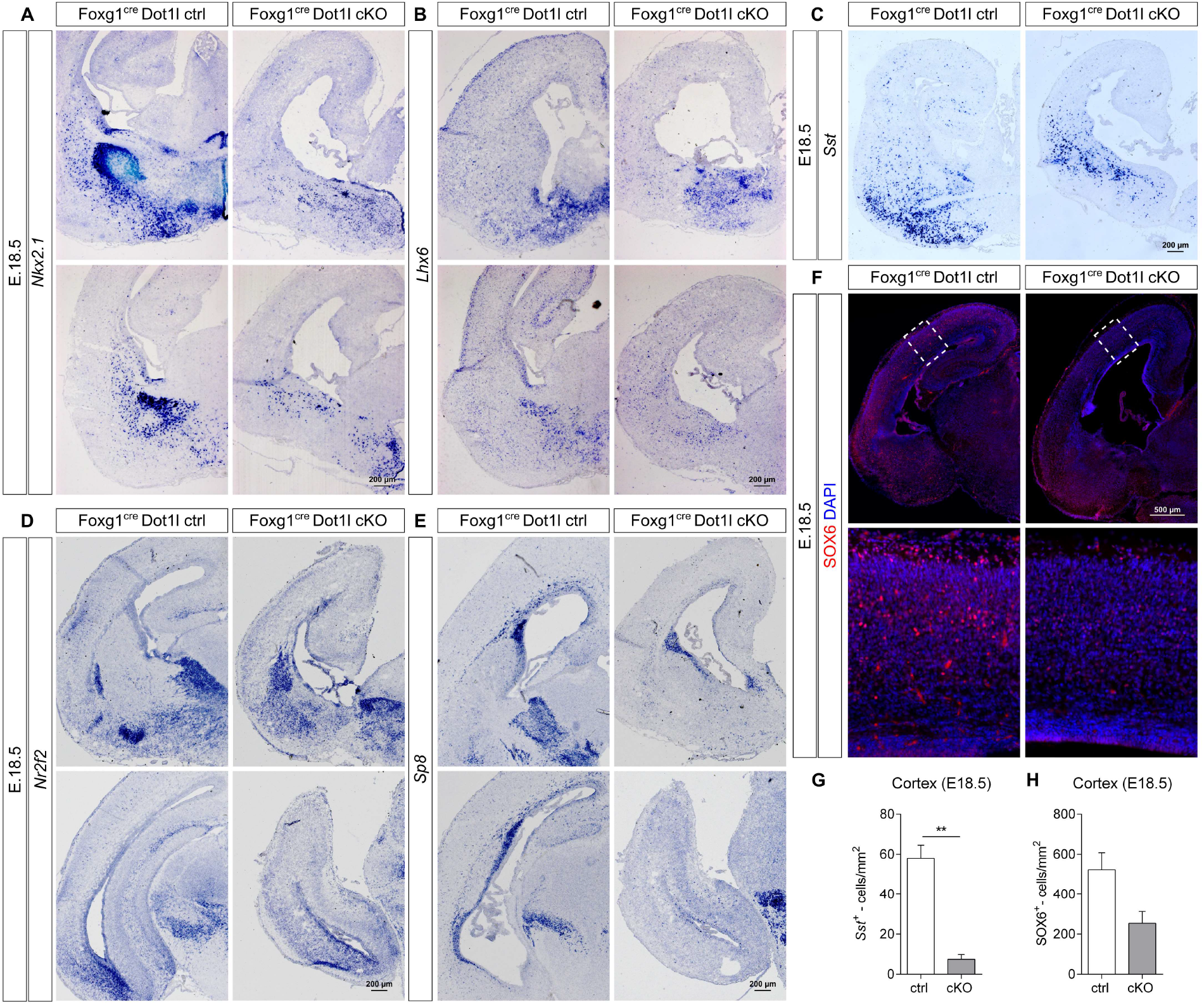
*Dot1l* is necessary for proper development of MGE- and CGE-derived interneurons. **(A-E)** Representative images of *in situ* hybridization (ISH) of embryonic brain sections using *Nkx2*.*1* (A), *Lhx6* (B), *Sst* (C), *Nr2f2* (D), and *Sp8* (E) antisense probes at embryonic stage E18.5. *Nkx2*.*1*, n=3-4; *Lhx6, Nr2f2* and *Sp8*, n=1 (Foxg1^cre^ Dot1l ctrl and cKO animals). Scale bars: 200 μm. **(F)** Representative images of immunostaining of E18.5 forebrain sections for SOX6 (red) and DAPI (blue); n=3 (Foxg1^cre^ Dot1l ctrl and cKO animals). Scale bars: 500 μm. **(G)** Quantification of *Sst*-expressing cells in the cerebral cortex of Foxg1^cre^ Dot1l cKO animals at E18.5. Data is represented as the mean ± SEM; n=3 (Foxg1^cre^ Dot1l ctrl and cKO animals); \*\**P* < 0.01 by unpaired, two tailed Student’s *t*-test with Welch’s correction. **(H)** Quantification of SOX6-expressing cells in the cortex of Foxg1^cre^ Dot1l cKO animals at E18.5. Data is represented as the mean ± SEM; n=3 (Foxg1^cre^ Dot1l ctrl and cKO animals); *P* > 0.05 by unpaired, two tailed Student’s *t*-test with Welch’s correction.

### *Dot1l* deletion in the Nkx2.1-cre lineage alters transcriptional programs of cortical interneurons

*Foxg1* is broadly expressed within both the ventral and dorsal telencephalon (Dou et al. 1999), and the concomitant *Dot1l* deficiency using Foxg1^cre^ could thus have cell autonomous and non-autonomous effects on the developing interneurons. In addition, Foxg1^cre^ Dot1l cKO mice die around birth, hampering detailed examination of interneuron classes that express relevant markers, e.g., PVALB, postnatally. Thus, to explore cell autonomous effects of DOT1L in the MGE interneuron lineage, we used Nkx2.1_cre_ as a driver line to generate *Dot1l*-deficient mice in the MGE lineage. For Nkx2.1^cre^ Dot1l cKO, we performed RNA sequencing for both ventral and dorsal telencephalon separately. RNA sequencing analysis of the ventral telencephalon from control (Nkx2.1^+/+^Dotl1^flox/+^) and cKO (Nkx2.1^cre/+^Dotl1^flox/flox^) mice at E14.5 revealed 747 DEGs (*p. adj. value* < 0.05), of which we classified 67 as interneuron marker genes (**Fig. 5A, B**). From the dorsal telencephalon we retrieved only 17 DEGs (*p. adj. value* < 0.1), of which only one classified as interneuron marker gene applying our curated short list (**Fig. S2A-C**) and two DEGs were shared between dorsal and ventral telencephalon data sets (**Fig. S2D**). Focusing on the 67 DEGs from the ventral telencephalon, we found both increased and decreased expression (**Fig. 5C**). Functional GO-term enrichment analysis revealed increased expression of genes associated to processes characteristic for post-mitotic neurons, such as regulation of membrane potential, neurotransmitter transport and synapse organization. Cell cycle-related terms, for instance cell cycle phase transition, DNA replication, or neural precursor cell proliferation, were enriched in genes with decreased expression (**Fig. 5D**). Accordingly, among DEGs with decreased expression upon *Dot1l* deletion, we observed enrichment of genes expressed in progenitor cells, whereas several markers for post-mitotic neurons showed increased expression (**Fig. S2E, F)**. Among the genes with decreased expression, we also observed the key regulatory transcription factors *Arx, Dlx1/2, Foxg1, Ldb1, Nkx2*.*1* and *Sp9* (**Fig. 5C**). Inspection of the H3K79me2 distribution at these genes rendered these interneuron targets as potential direct DOT1L regulated genes (**Fig. 3B, C, 5E**). Compared to the Foxg1^cre^ Dot1l cKO we identified 162 common target genes with the Nkx2.1^cre^ mediated cKO, 16 of which classified as interneuron markers (**Fig. S2G, H**). Shared DEGs between both mouse lines contained a set of genes with opposing expression changes, but around 2/3 of DEGs decreased in both models. Compromised expression of instructors of interneuron development affected in both cKO models included *Arx, Lmo1, Sox1/2, Tcf4* and *Zswim5* (**Fig. S2H**). Together, our data indicated that DOT1L is implicated in proper embryonic development of MGE-derived interneurons of the *Nkx2*.*1* lineage by regulating expression of key transcription factors.

**Figure 5.**
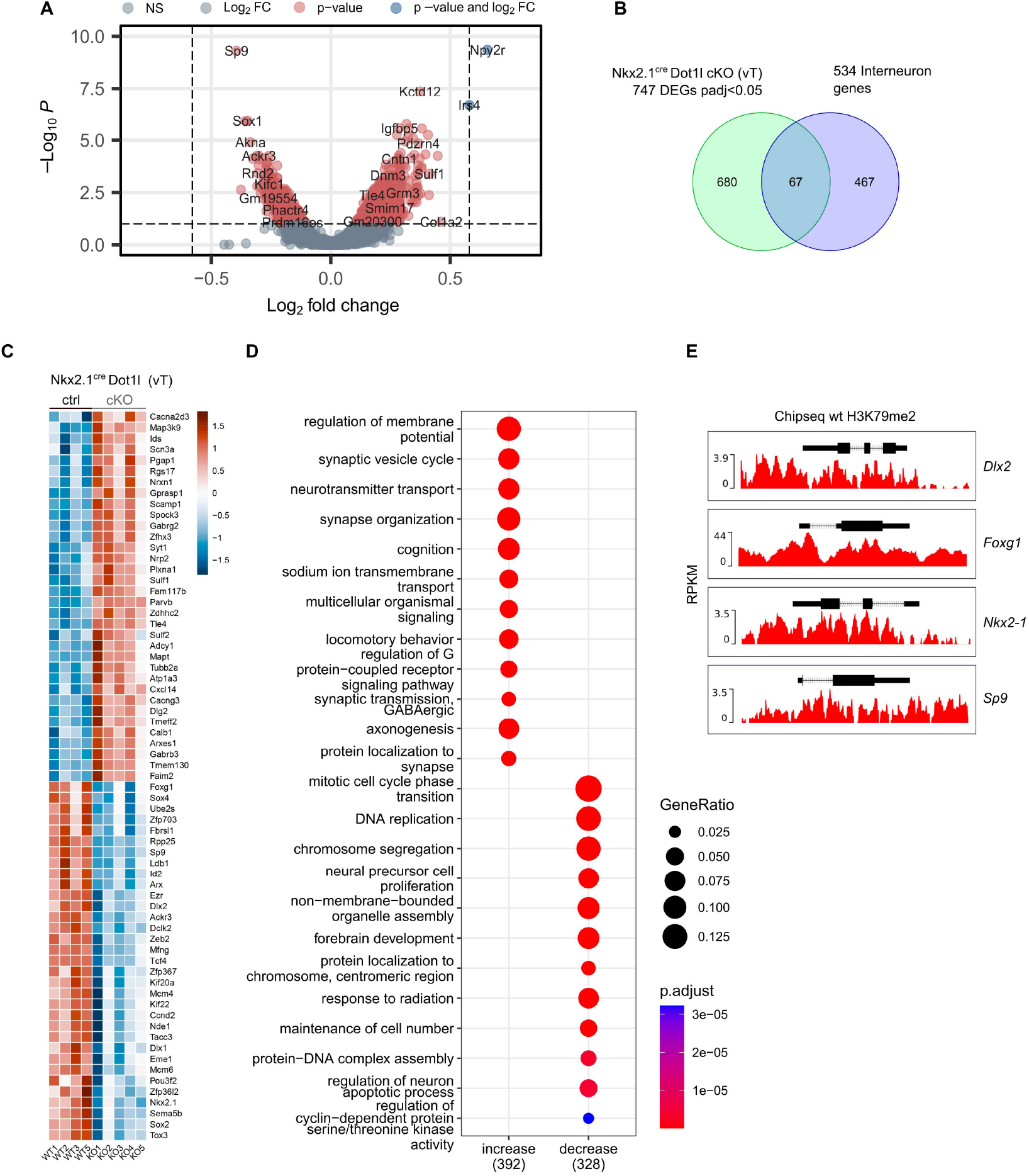
*Dot1l* deletion in the Nkx2.1-cre lineage impairs cortical interneuron development. **(A)** Volcano plot displaying differentially expressed genes (DEGs) in the ventral telencephalon of Nkx2.1^cre^ Dot1l cKO animals compared to controls at E14.5. The y-axis corresponds to the adjusted p-value and the x-axis displays the log_2_Fold Change (log_2_FC). Grey: Transcripts with insignificant adjusted p-values (p > 0.05). Red: Transcripts with differential expression of less than ± log_2_ (0.5). Blue: Transcripts with differential expression of more than ± log_2_ (0.5). Positive log_2_FC represents increase, negative log_2_FC represents decrease in expression upon *Dot1l* deletion. **(B)** Venn diagram showing the intersection of DEGs in ventral telencephalon (vT) of Nkx2.1^cre^ Dot1l cKO animals at E14.5 and genes classified as interneuron marker genes. **(C)** Heatmap of 67 intersected genes in (B). DEGs with increased or decreased expression are represented in red or blue, respectively. Scale shows scaled VST normalized gene counts. Data represents sex- and batch-corrected gene expression in vT of Nkx2.1^cre^ Dot1l control (n=4) and cKO (n=5) animals. **(D)** Dotplot of differential GO term enrichment analysis of increasing and decreasing DEGs in Nkx2.1^cre^ Dot1l cKO mice. Gene ratios and adjusted p-values are indicated at the right and total number of genes per group is shown on the x-axis. Blue depicts the least significantly enriched pathway, and red the most significantly enriched pathway. **(E)** ChIP-seq results of H3K79me2 levels at *Dlx2, Foxg1, Nkx2*.*1* and *Sp9* genes in E14.5 wildtype cerebral cortex, a set of candidate genes, which also decreased in expression in the ventral telencephalon in Nkx2.1^cre^ Dot1l cKO animals at E14.5 as revealed by RNA-seq and shown in (C). Red peaks indicate enrichment for H3K79me2 of genes with decreased expression.

### *Dot1l* deletion in the Nkx2.1-cre lineage reduces the number of cortical interneurons

We next explored expression of interneuron markers by histological methods *in vivo*. ISH at E14.5 and E18.5 revealed reduced expression of *Nkx2*.*1* (**Fig. 6A, C**). As expected, the *Nkx2*.*1* downstream effector *Lhx6* (Sandberg et al. 2016) also decreased in its expression at both developmental time points (**Fig. 6B, D**). The CGE markers *Nr2f2* and *Prox1* (both at E14.5 and E18.5) and the dorsal MGE marker *Nkx6*.*2* (at E14.5) did show neither differential expression in the transcriptome, nor striking different staining patterns using ISH, supporting the specificity of the cKO in the MGE lineage (**Fig. S3A-E**).

**Figure 6.**
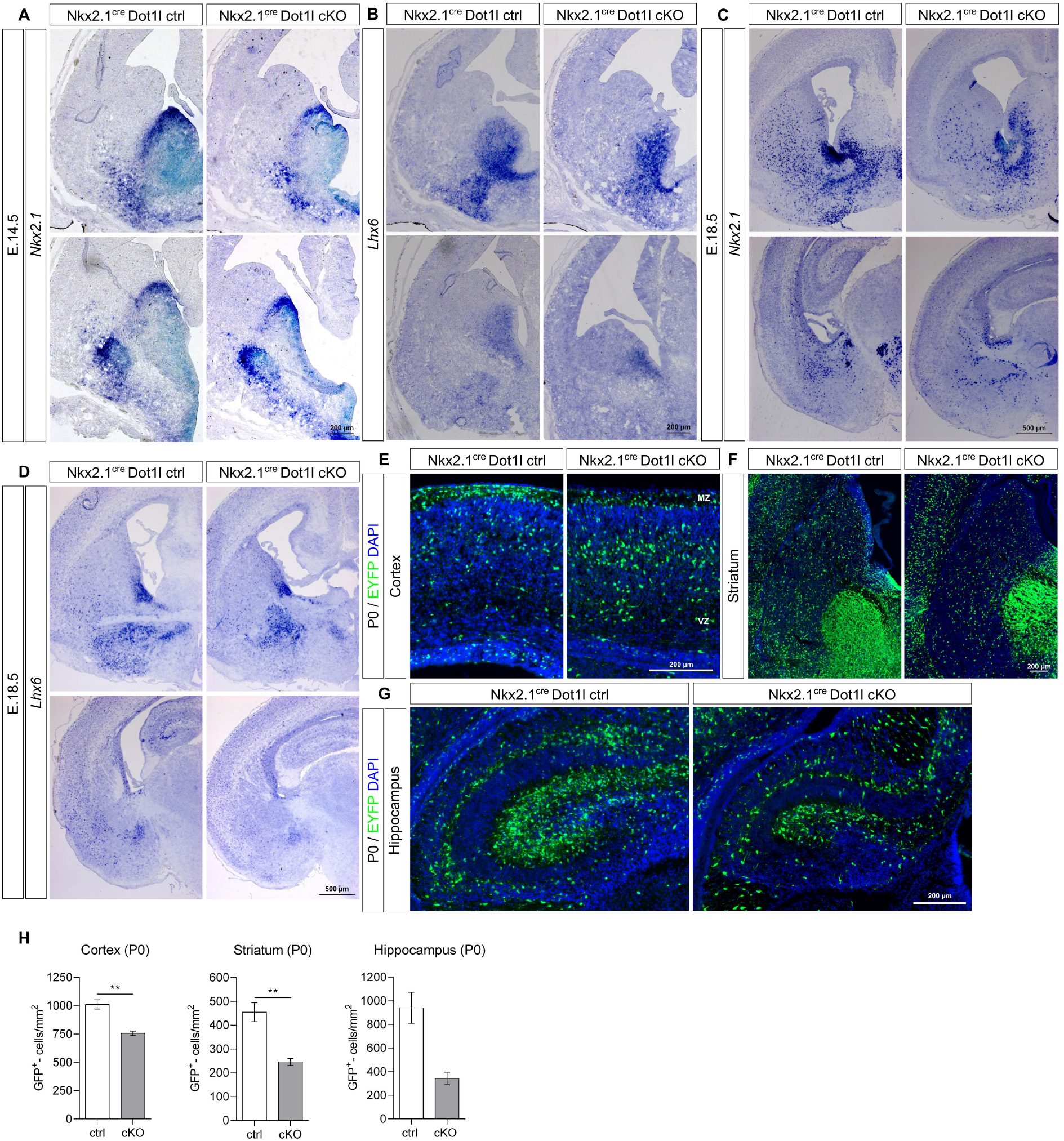
*Dot1l* deletion in the Nkx2.1-cre lineage reduces the number of cortical interneurons at different developmental stages. **(A-D)** Representative images of *in situ* hybridization using *Nkx2*.*1* (A) and *Lhx6* (B) antisense probes at E14.5, and E18.5 (C, D), showing lower expression of these MGE markers; n=1. Scale bars: 200 μm (A, B) and 500 μm (C, D). **(E-G)** Representative images of immunostaining of postnatal (P0) forebrain sections for GFP (EYFP) (green) and DAPI (blue) in the cerebral cortex (A), hippocampus (B) and striatum (C), n=2-4 (Nkx2.1^cre^ Dot1l control and cKO animals). Scale bars: 200 μm. **(H)** Quantification of MGE-derived GABAergic interneurons (GFP^+^ cells) in the postnatal cortex, striatum and hippocampus. Data is represented as the mean ± SEM; Cortex and striatum: n=4 (Nkx2.1^cre^ Dot1l ctrl and cKO animals); Hippocampus: n=2-3 (Nkx2.1^cre^ Dot1l ctrl and cKO animals); \*\**P* < 0.01 by unpaired, two tailed Student’s *t*-test with Welch’s correction.

For gaining insight into how the observed transcriptional changes upon DOT1L deletion would impact MGE-derived interneurons at postnatal stages, we characterized these interneurons at postnatal day 0 (P0), when the interneurons completed their migration to the cerebral cortex, and at P21, when lamination as well as circuit integration and refinement are completed. To this end, we traced the MGE-derived interneurons using a ROSA26-YFP reporter mouse line crossed with the Nkx2.1^cre^ Dotl1 line. A statistically significant decrease in the number of MGE-derived interneurons was observed in the cortex (757.10 ± 18.03 cells/mm^2^ *versus* 1011 ± 39.64 cells/mm^2^; \*\**p* < 0.01) and striatum (246.00 ± 14.72 cells/mm^2^ *versus* 457.70 ± 40.14 cells/mm^2^; \*\**p* < 0.01) of cKO mice at P0 compared to control animals (**Fig. 6E, F, H)**. In the hippocampus, cell numbers were non-significantly decreased (343.50 ± 52.62 cells/mm^2^ *versus* 942.00 ± 131.70 cells/mm^2^; *p* > 0.05) (**Fig. 6G, H**).

Both, somatostatin (SST) and parvalbumin (PVALB)-expressing interneurons are born in the MGE, and we next explored whether deletion of *Dot1l* in *Nkx2*.*1*-expressing precursors would affect both lineages or have lineage-specific effects. At P0, we quantified only the number of the *Sst*-expressing subfraction of interneurons, since PVALB expression is restricted to later postnatal stages. *Dot1l* cKO mice possibly exhibited fewer *Sst*-positive interneurons, however, our quantification did not reach statistical significance (**Fig. S4**).

At P21, we performed single molecule (sm)FISH for *Pvalb* and *Sst* as well as for *Npy*, an interneuron marker usually co-localized with *Sst*, to further discriminate between the MGE-derived interneuron populations. The total number of MGE-derived interneurons, i.e. *Pvalb, Sst* and *Npy/Sst* expressing cells, in the cerebral cortex of cKO was significantly reduced (55.64 ± 7.67 cells/mm^2^ *versus* 107 ± 7.907 cells/mm^2^; \*\**p* < 0.01), while MGE-derived interneuron counts in the striatum showed only a trend towards decreased numbers in cKO mice compared to controls (**Fig. 7A, B**). *Pvalb*-expressing interneurons decreased significantly in both cortex and striatum of cKOs compared to controls (**Fig. 7C**), but both *Sst* and *Sst/Npy* did not change (**Fig. 7D, E**).

**Figure 7.**
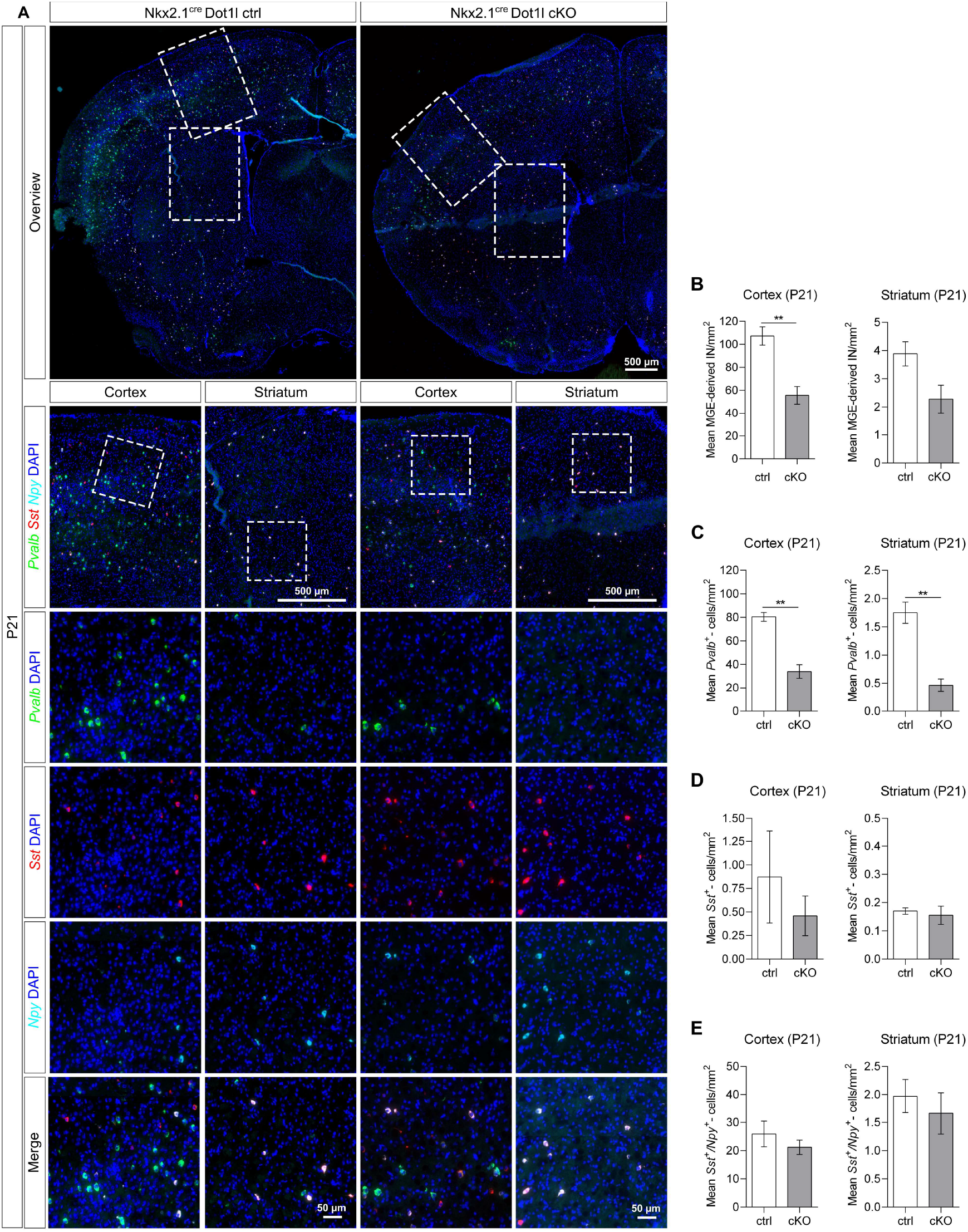
*Dot1l* deletion in the Nkx2.1-cre lineage reduces the number of cortical *Pvalb*-expressing interneurons at postnatal day 21 (P21) **(A)** Representative images of parvalbumin (*Pvalb*), somatostatin (*Sst*) and neuropeptide-Y (*Npy*) expressing MGE-derived interneurons using single molecule FISH at P21; n=3-4 (Nkx2.1^cre^ Dot1l ctrl and cKO animals). Scale bar: 500 μm. Higher magnification of representative images of cerebral cortex and striatum regions, and higher magnifications of individual *Pvalb* (green), *Sst* (red) or *Npy* (cyan) expressing GABAergic interneurons together with DAPI (blue), and the combination of all (merge). Scale bar: 50 μm. **(B)** Quantification of total MGE-derived interneurons (*Pvalb* (C), *Sst* (D) and *Sst/Npy* (E)) in the cortex and striatum of Nkx2.1^cre^ Dot1l cKO animals compared to controls at P21. Data is represented as the mean ± SEM; n=3-4 (Nkx2.1^cre^ Dot1l ctrl and cKO animals); \*\**P* < 0.01 by unpaired, two tailed Student’s *t*-test with Welch’s correction. **(C-E)** Quantification of *Pvalb* (C), *Sst* (D) and *Sst/Npy* (E) expressing interneurons in the cerebral cortex and striatum of Nkx2.1^cre^ Dot1l cKO animals compared to controls at P21. Data is represented as the mean ± SEM; n=3-4 (Nkx2.1^cre^ Dot1l ctrl and cKO animals); \*\**P* < 0.01 by unpaired, two tailed Student’s *t*-test with Welch’s correction.

Since PVALB expression is not detectable at embryonic and early postnatal stages, we used two markers for immature *Pvalb-*expressing interneurons, *Phlda1* and *Plcxd3* (Mayer et al. 2018), to assess whether immature *Pvalb* interneurons were reduced at the embryonic development stages E14.5 and E16.5, and at the postnatal stage P0 (**Fig. S5A**). We did not observe statistically significant changes in cell numbers for *Phlda1* and *Plcxd3* expression.

We further assessed whether the reduction of MGE-derived interneurons, most likely affecting the *Pvalb*-expressing fraction, upon *Dot1l* cKO, would be caused by increased apoptosis and/or reduced proliferation of MGE progenitors at E14.5. We did neither observe increased numbers of cells expressing aCASP3 in cKO mice, nor did we observe significant changes of cycling BrdU-labeled progenitors (**Fig. S5B-D**). These experiments made it unlikely that apoptosis or decreased proliferation of MGE progenitors upon *Dot1l* cKO contributed to the decreased numbers of interneurons.

Taking all data together, we conclude that DOT1L affects the MGE-derived PVALB subpopulation of interneurons by regulating the specification, maturation and/or migration.

## Discussion

In this work, we used two different cKO mouse models, namely Foxg1^cre^ Dot1l and Nkx2.1^cre^ Dot1l cKO lines and demonstrated that the epigenetic modifier DOT1L affects embryonic and postnatal development of interneurons. *Dot1l* deletion in forebrain progenitors through Foxg1^cre^ affected transcriptional programs in both MGE- and CGE-derived interneurons. *Dot1l* deletion in the MGE through Nkx2.1^cre^ resulted in reduced numbers of cells expressing *Nkx2*.*1* and *Lhx6* in cKO mice compared to control animals. Specifically, loss of *Dot1l* in MGE-located progenitors impaired the *Pvalb*-expressing subpopulation of interneurons, which reduced in numbers postnatally. Moreover, upon loss of *Dot1l*, genes related to cell cycle decreased, while the expression of neuronal markers increased. This finding strongly pointed towards neuronal differentiation or specification/maturation defects as possible underlying mechanisms.

That *Dot1l* deletion in *Foxg1*-expressing cells had broader effects on the interneuron development regarding the affected populations might reflect the fact that *Foxg1* has a wider expression in the ganglionic eminences, being expressed by both MGE- and CGE-derived cells (Dou et al. 1999). For the DEGs shared by our two models we also observed that the transcriptional changes were usually stronger for the Foxg1^cre^ Dot1l cKO, when compared to the Nkx2.1^cre^ Dot1l cKO mouse model. A reason for this difference might be that the bulk RNAseq data for Foxg1^cre^ Dot1l control and cKO mice were obtained from the dorsal telencephalon, and that the *Dot1l* deletion in this model was not restricted to GABAergic progenitors alone, but also encompassed glutamatergic progenitors. This loss of DOT1L in both progenitor populations adds another source impacting interneuron development, namely, the environment sensed by the cells and cell-to-cell interactions (Achim et al. 2014; Brandão and Romcy-Pereira 2015). The microenvironment might be differentially shaped in Foxg1^cre^*-*mediated *Dot1l*, and the interneurons devoid of the epigenetic modifier entered a mis-specified target region. It is thus tempting to speculate, but beyond the scope of this paper, that restricted deletion of *Dot1l* in the cortical plate will impact interneuron development as well.

*Dot1l* deletion in MGE progenitor cells interfered specially with the development of *Pvalb*-expressing interneurons. We cannot fully explain this observation based on our data as of yet. First, we cannot completely rule out that decreased proliferation might contribute for the reduction of *Pvalb*-expressing interneurons, although cell cycle analysis using histological methods revealed only a tendency towards decreased proliferation and increased cell cycle exit upon *Dot1l* cKO in the MGE (**Fig. S6C**). We provided strong evidence in other parts of the CNS that DOT1L is crucial for maintaining the progenitor pool and consequently suitable neuronal numbers. For example either pharmacological inhibition or short hairpin RNA (shRNA)-mediated knockdown of DOT1L results in decreased proliferation of primary cortical neural stem cells (Roidl et al. 2016). Similar results were recapitulated by *in vitro* neuronal differentiation of NPCs derived from mouse embryonic stem cells (mESCs) upon pharmacological inhibition of DOT1L, where DOT1L keeps the NPCs in a proliferative state through a mechanism dependent on the accessibility of SOX2-bound enhancers (Ferrari et al. 2020). Moreover, genetic deletion of *Dot1l* in the glutamatergic lineage led to increased cell cycle exit (Franz et al. 2019). Since *Pvalb* interneurons are born later than the *Sst* expressing subpopulations (Wonders et al. 2008; Inan et al. 2012), it is tempting to speculate that the reduced numbers of *Pvalb* interneurons are caused by a mild, but not statistically significant reduction of the progenitor pool, that only becomes noticeable in the last subpopulation generated, i.e., *Pvalb-*expressing interneurons.

Second, in addition to temporal differences, spatial information could contribute to the *Pvalb*-specific effect of *Dot1l* deletion on interneuron development. The dorsal part from the MGE produces mostly SST-positive interneurons, and progenitors in the ventral MGE give rise to PVALB-positive interneurons (Wonders et al. 2008). It is thus tempting to speculate that *Dot1l* deletion influences differently the cell fate decisions in the dorsal and ventral MGE, with the ventral MGE being more affected than the dorsal MGE. However, we did not observe changes in the expression of the dorsal MGE marker *Nkx6*.*2* (**Fig. S3C**).

By using BrdU birthdating experiments combined with transplantation studies, different authors have shown that, while SST-expressing interneurons are mostly generated at earlier time points (E12.5/13.5), PVALB-expressing interneurons are continuously born between E12.5 and E15.5 (Wonders et al. 2008; Inan et al. 2012). This is in agreement with the observation that fate specification of interneurons is also related to the location of neurogenesis in the MGE. SST-expressing interneurons are mostly generated from apical progenitors in the ventricular zone, whereas late CCND2-expressing basal progenitors in the subventricular zone mostly give rise to PVALB-expressing interneurons (Petros et al. 2015). *Ccnd2* was one target gene with decreased expression upon *Dot1l* deletion in the MGE (**Fig. 5C**). We therefore hypothesize that *Dot1l* cKO-induced reduction in the number of *Pvalb*-expressing interneurons is due to a reduction of the basal progenitor pool, which results from a mild reduced proliferation of apical progenitors and their premature cell cycle exit followed by differentiation into *Sst*-expressing interneurons before the transition to basal progenitors. This scenario would explain the discrete tendency to decreased numbers of *Sst*-expressing interneurons, while statistical significance for *Pvalb*-expressing interneurons was reached. In favor of this interpretation is our earlier report that upon *Dot1l* cKO, cortical progenitor cells exit the cell cycle prematurely and adapt DL neuron fate. At the time of the transition from apical to basal progenitors, there are less apical and consequently less TBR2-positive basal progenitors to give rise to UL neurons (Franz et al. 2019). Supporting our hypothesis is also the recent observation that the cortical interneuron populations result from a post-mitotic lineage divergence. Interneurons derive from a common progenitor pool with limited heterogeneity, and, soon after cell cycle exit, different transcriptional programs refine the trajectories taken by the post-mitotic interneurons and, consequently, their identities (Bandler et al. 2022).

Additionally, we observed an aberrant lamination of cortical glutamatergic DL neurons (Franz et al. 2019), which might also parallel some of our observations regarding the mis-localization of MGE-derived interneurons upon *Dot1l* cKO that mainly populated superficial layers of the cortical plate, i.e., layers 4-6 **(Fig. 6E**). PVALB-positive neurons settle preferentially between layers 1 and 2, and we observed fewer neurons in these layers upon loss of DOT1L (Miyamae et al. 2017). The mis-localization of interneurons might thus reflect an accelerated or premature neuronal differentiation and subsequently, earlier invasion of the interneurons in the cortical plate concomitant with a failure to adapt a mature PVALB fate/localization. Taking into account the observed decreased expression of genes related to cell cycle, including the basal progenitor marker and the increased activity of genes involved in late neuronal maturation (**Fig. 7**), the maintenance of the progenitor pool by DOT1L might also play an important role in the development of MGE-derived interneurons, as we have observed for cortical glutamatergic neurons.

The altered distribution of MGE-derived interneurons in the cortical plate at P0 upon *Dot1l* deletion (**Fig. 5**), might not only indicate premature differentiation and invasion of the cortical plate, but also impaired post-mitotic maturation. One possible mechanism might be attributed to the decreased expression of *Ackr3* in the ventral telencephalon from Nkx2.1^cre^ Dotl1 cKO mice at E14.5 (**Fig. 1C** and **4B**). ACKR3 (also known as CXCR7) is a CXCL12 receptor expressed by migrating interneurons and this chemokine signaling pathway is important for proper interneuron migration (López-Bendito et al. 2008; Bartolini et al. 2017; Sánchez-Alcañiz et al. 2011). Interneurons of *Ackr3*^-/-^ mice leave the MZ and the SVZ to accumulate in the cortical plate. Thus, one might hypothesize that the reduced expression of *Ackr3* upon *Dot1l* deletion in MGE-derived interneurons is involved in their aberrant laminar distribution in the cortical plate.

Additional research to address open, mechanistic questions will still be needed to increase our understanding on the epigenetics and specifically DOT1L functions governing the GABAergic development in normal and disease conditions.

## Acknowledgments

We thank C. Fullio for her help plotting data with ‘R’ and S. Heinrich for her help with animal work. This work was supported by the Deutsche Forschungsgemeinschaft (DFG) through a grant to TV (VO1676/7-1). Further, the authors thank the Freiburg Galaxy Server team and all members of the Vogel group for discussion and support.

