## Supplementary figures and tables for "DOT1L deletion impairs the development of cortical Parvalbumin-expressing interneurons"

### Figure S1

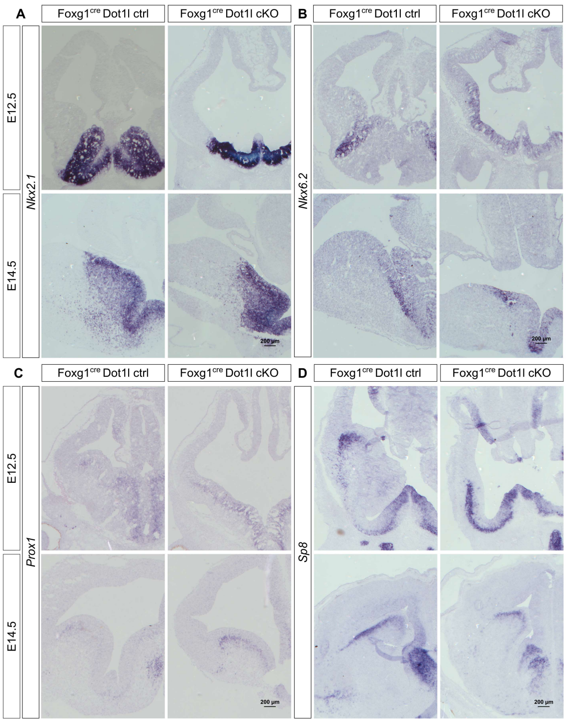

A
Foxg1cre Dot1l ctrl
Foxg1cre Dot1l cKO
B
Foxg1cre Dot1l ctrl
Foxg1cre Dot1l cKO
E12.5
E14.5
Nkx2.1
Nkx6.2
200 µm
200 µm
C
Foxg1cre Dot1l ctrl
Foxg1cre Dot1l cKO
D
Foxg1cre Dot1l ctrl
Foxg1cre Dot1l cKO
E12.5
E14.5
Prox1
Sp8
200 µm
200 µm

### Figure S2

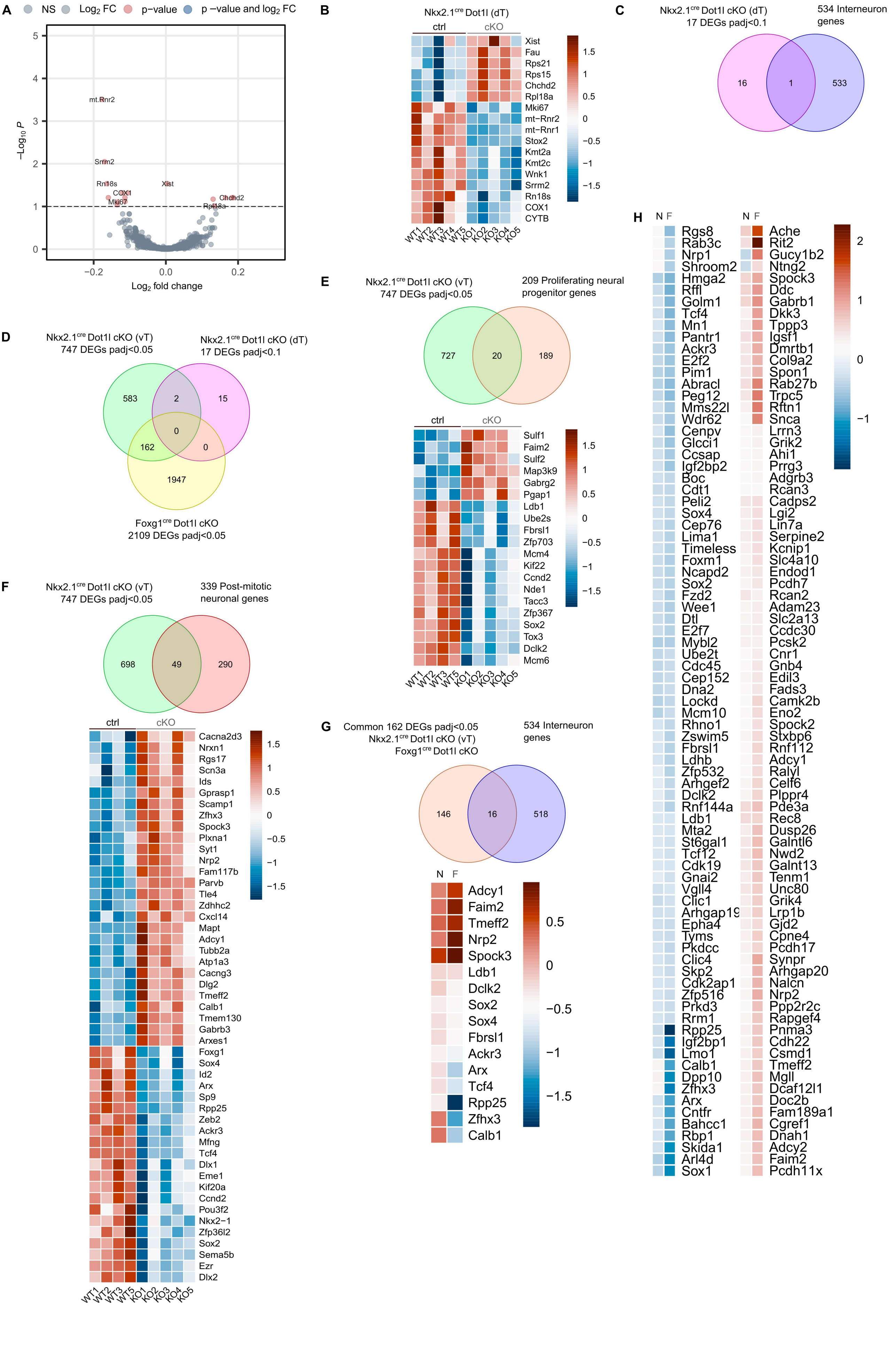
A
NS
Log2 FC
p−value
p −value and log2 FC
−Log10 P
Log2 fold change
mt.Rnr2
Srrm2
Rn18s
COX1
Mki67
Xist
Chchd2
Rpl18a
B
Nkx2.1cre Dot1l (dT)
ctrl
cKO
Xist
Fau
Rps21
Rps15
Chchd2
Rpl18a
Mki67
mt−Rnr2
mt−Rnr1
Stox2
Kmt2a
Kmt2c
Wnk1
Srrm2
Rn18s
COX1
CYTB
WT1 WT2 WT3 WT4 WT5 KO1 KO2 KO3 KO4 KO5
C
Nkx2.1cre Dot1l cKO (dT)
17 DEGs padj<0.1
534 Interneuron genes
16
1
533
D
Nkx2.1cre Dot1l cKO (vT)
747 DEGs padj<0.05
Nkx2.1cre Dot1l cKO (dT)
17 DEGs padj<0.1
583
2
15
0
162
0
1947
Foxg1cre Dot1l cKO
2109 DEGs padj<0.05
E
Nkx2.1cre Dot1l cKO (vT)
747 DEGs padj<0.05
209 Proliferating neural progenitor genes
727
20
189
ctrl
cKO
Sulf1
Faim2
Sulf2
Map3k9
Gabrg2
Pgap1
Ldb1
Ube2s
Fbrsl1
Zfp703
Mcm4
Kif22
Ccnd2
Nde1
Tacc3
Zfp367
Sox2
Tox3
Dclk2
Mcm6
WT1 WT2 WT3 WT5 KO1 KO2 KO3 KO4 KO5
F
Nkx2.1cre Dot1l cKO (vT)
747 DEGs padj<0.05
339 Post-mitotic neuronal genes
698
49
290
ctrl
cKO
Cacna2d3
Nrxn1
Rgs17
Scn3a
Ids
Gprasp1
Scamp1
Zfhx3
Spock3
Plxna1
Syt1
Nrp2
Fam117b
Parvb
Tle4
Zdhhc2
Cxcl14
Mapt
Adcy1
Tubb2a
Atp1a3
Cacng3
Dlg2
Tmeff2
Calb1
Tmem130
Gabrb3
Arxes1
Foxg1
Sox4
Id2
Arx
Sp9
Rpp25
Zeb2
Ackr3
Mfng
Tcf4
Dlx1
Eme1
Kif20a
Ccnd2
Pou3f2
Nkx2−1
Zfp36l2
Sox2
Sema5b
Ezr
Dlx2
WT1 WT2 WT3 WT5 KO1 KO2 KO3 KO4 KO5
G
Common 162 DEGs padj<0.05
Nkx2.1cre Dot1l cKO (vT)
Foxg1cre Dot1l cKO
534 Interneuron genes
146
16
518
N F
Adcy1
Faim2
Tmeff2
Nrp2
Spock3
Ldb1
Dclk2
Sox2
Sox4
Fbrsl1
Ackr3
Arx
Tcf4
Rpp25
Zfhx3
Calb1
H
N F
Rgs8
Rab3c
Nrp1
Shroom2
Hmga2
Rffl
Golm1
Tcf4
Mn1
Pantr1
Ackr3
E2f2
Pim1
Abracl
Peg12
Mms22l
Wdr62
Cenpv
Glcci1
Ccsap
Igf2bp2
Boc
Cdt1
Peli2
Sox4
Cep76
Lima1
Timeless
Foxm1
Ncapd2
Sox2
Fzd2
Wee1
Dtl
E2f7
Mybl2
Ube2t
Cdc45
Cep152
Dna2
Lockd
Mcm10
Rhno1
Zswim5
Fbrsl1
Ldhb
Zfp532
Arhgef2
Dclk2
Rnf144a
Ldb1
Mta2
St6gal1
Tcf12
Cdk19
Gnai2
Vgll4
Clic1
Arhgap19
Epha4
Tyms
Pkdcc
Clic4
Skp2
Cdk2ap1
Zfp516
Prkd3
Rrm1
Rpp25
Igf2bp1
Lmo1
Calb1
Dpp10
Zfhx3
Arx
Cntfr
Bahcc1
Rbp1
Skida1
Arl4d
Sox1
N F
Ache
Rit2
Gucy1b2
Ntng2
Spock3
Ddc
Gabrb1
Dkk3
Tppp3
Igsf1
Dmrtb1
Col9a2
Spon1
Rab27b
Trpc5
Rftn1
Snca
Lrrn3
Grik2
Ahi1
Prrg3
Adgrb3
Rcan3
Cadps2
Lgi2
Lin7a
Serpine2
Kcnip1
Slc4a10
Endod1
Pcdh7
Rcan2
Adam23
Slc2a13
Ccdc30
Pcsk2
Cnr1
Gnb4
Edil3
Fads3
Camk2b
Eno2
Spock2
Stxbp6
Rnf112
Adcy1
Ralyl
Celf6
Plppr4
Pde3a
Rec8
Dusp26
Galntl6
Nwd2
Galnt13
Tenm1
Unc80
Grik4
Lrp1b
Gjd2
Cpne4
Pcdh17
Synpr
Arhgap20
Nalcn
Nrp2
Ppp2r2c
Rapgef4
Pnma3
Cdh22
Csmd1
Tmeff2
Mgll
Dcaf12l1
Doc2b
Fam189a1
Cgref1
Dnah1
Adcy2
Faim2
Pcdh11x

### Figure S3

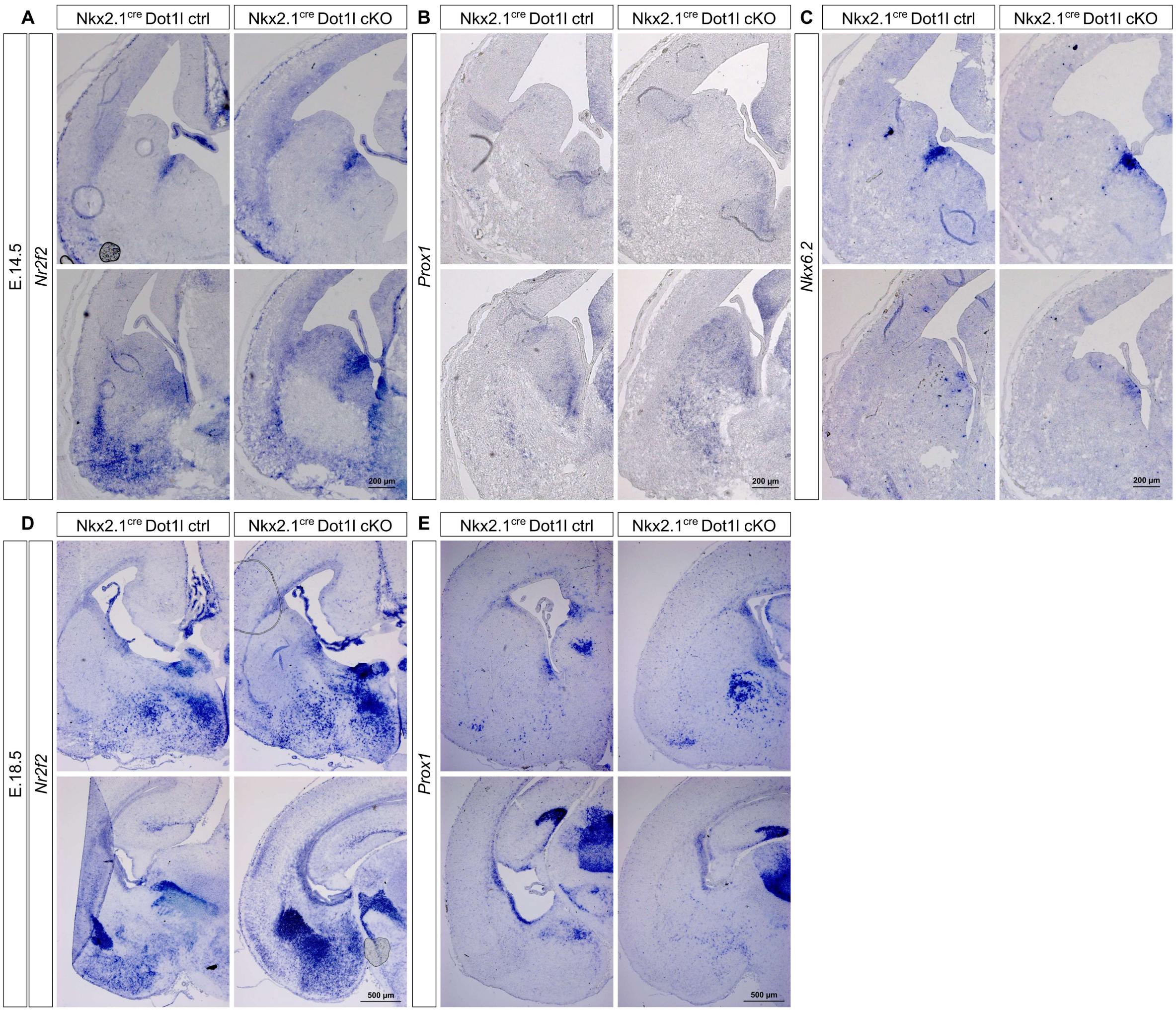

A
B
C
D
E
Nkx2.1cre Dot1l ctrl
Nkx2.1cre Dot1l cKO
E.14.5
E.18.5
Nr2f2
Prox1
Nkx6.2
200 µm
500 µm

### Figure S4

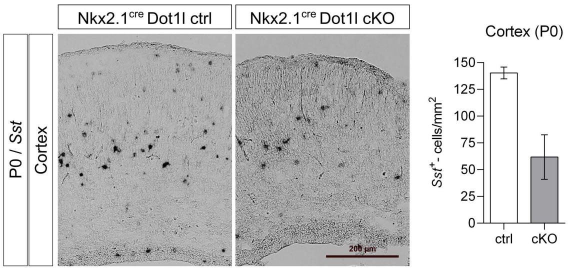
Nkx2.1cre Dot1l ctrl
Nkx2.1cre Dot1l cKO
P0 / Sst
Cortex
200 µm
Cortex (P0)
Sst+ - cells/mm2
150
125
100
75
50
25
0
ctrl
cKO

### Figure S5

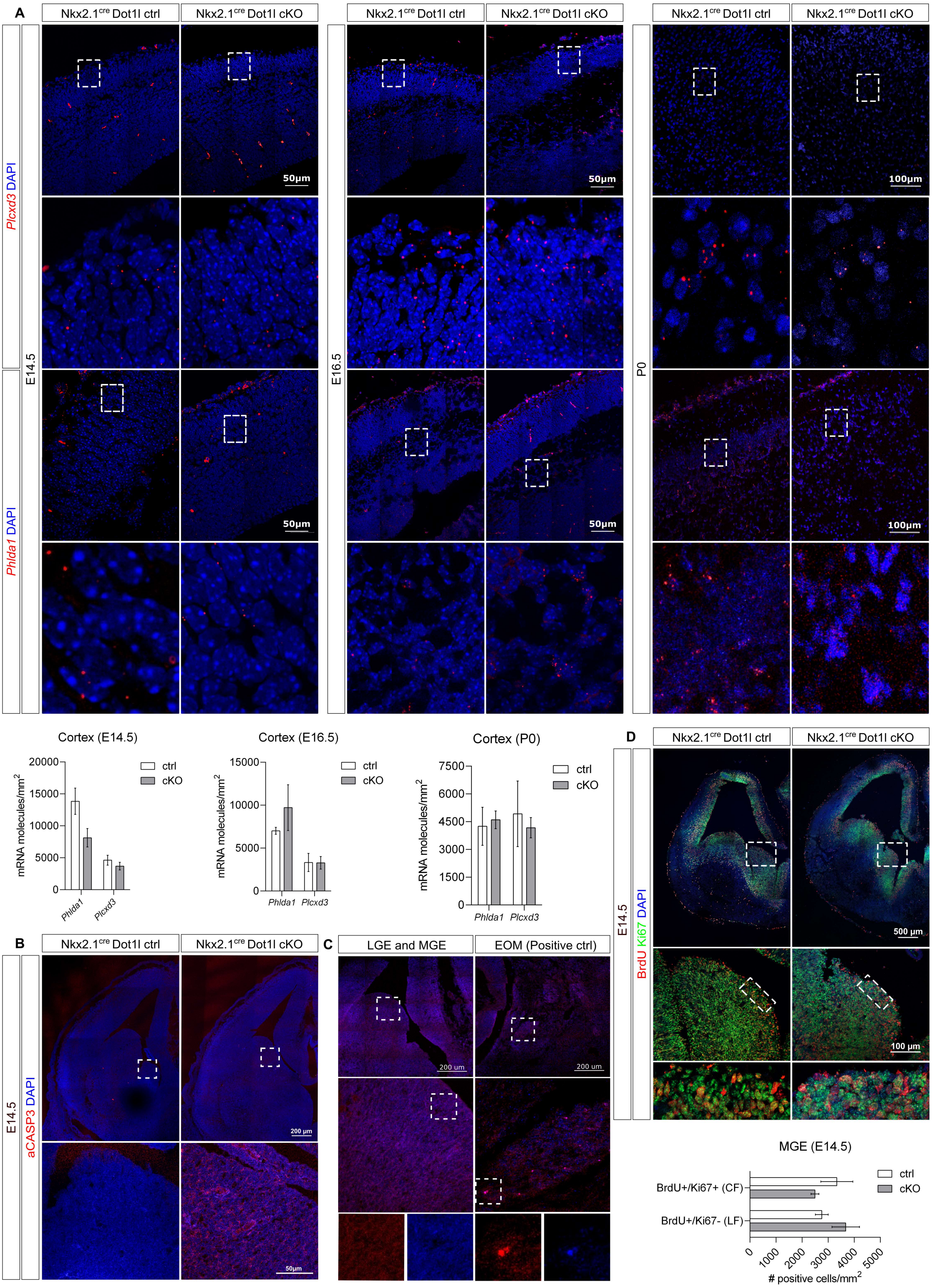
A
Nkx2.1cre Dot1l ctrl
Nkx2.1cre Dot1l cKO
Plcxd3 DAPI
Phlda1 DAPI
E14.5
E16.5
P0
50μm
100μm
Cortex (E14.5)
Cortex (E16.5)
Cortex (P0)
mRNA molecules/mm2
ctrl
cKO
Phlda1
Plcxd3
B
C
LGE and MGE
EOM (Positive ctrl)
aCASP3 DAPI
200 μm
200 um
D
BrdU Ki67 DAPI
500 μm
100 μm
MGE (E14.5)
BrdU+/Ki67+ (CF)
BrdU+/Ki67- (LF)
### positive cells/mm2

**Supplementary Table 1** List of qRTPCR primers

| **Target region** | **Reverse Primer** | **Forward Primer** |
|---|---|---|
| *Ache* | CCAGAGTATCGGTGGCGCT | GACCAATTTTGCCCGCACAG |
| *Arx* | CAGCTCAGCCTCGAACGG | TCCAACCCTCCAGGAGAGAG |
| *Ascl1* | CCCATCTGTGATTCGGGCTT | CAGACAGCCAACCTACAGGG |
| *Calb1* | GTGGGTAAGACGTGAGCCAA | ATTTCGACGCTGACGGAAGT |
| *Calb2* | GGCCAAGGACATGACACTCT | GATGAGAATGAACTGGACGCC |
| *Ccnd1* | GGAAGACCTCCTCTTCGCAC | GAGATTGTGCCATCCATGCG |
| *Ccnd2* | GCCAAGAAACGGTCCAGGTA | TACCTCCCGCAGTGTTCCTA |
| *Chrna7* | ATGCCCAGATGCATTCACCA | ACATGTCTGAGTACCCCGGA |
| *Cux2* | CTGAACACCGCCGTACTCC | CGGACATCCATCTCTTAGCCC |
| *Dot1l* | GCATCATGGTGCTTGTCGTA | TGAGGCTCAAGTCGCCTGT |
| *Dlx1* | AACTTGGAGCGTTTGTTCTGG | CGGAGGTTCCAACAAACTCAG |
| *Dlx2* | ACTGGAGTAGATGGTGCGTG | AGTTCGTCTCCGGTCAACAA |
| *Dlx5* | GCCCATCTAATAAAGCGTCCC | CTACCTGGAGAACTCGGCTT |
| *Dlx6* | GGTCACTCTCGTGTGGGTT | CAGACTCAATACCTGGCCCT |
| *Erbb4* | TGGTAAAGTGGAATGGCCCG | CAGATCAGGATCGGGAGTGC |
| *Htr3a* | GGTGATCGCCACAGGGTAAT | ACAAGCCCTTGCAATTGGTG |
| *Ldb1* | GTTGGCTCCACCTTAGTCCT | GCTCCTTCGGCGAGTACAG |
| *Lhx5* | ATCTCAGACTCGTCCAAGCG | TGAGGGTCATTCAGGTGTGG |
| *Lhx6* | TTTGCAGCCGGACCCTG | TGGAAGCATGAGAGCGCC |
| *Lhx9* | CGCATCCGTTTGGTCTTCTG | CTGGGAGTGGACATCGTGAA |
| *Lmo1* | GGCGAAGCAGTCAAGGTGAT | AAGGCCAACCTCATCCTGTG |
| *Maf* | GGTCTCCACCGGTTCCTTTT | AGGATGGCTTCAGAACTGGC |
| *Mid1* | AGCAAAGTCCAAGGAAGCTGA | GGAGGGGGACGAAAAACGTG |
| *Nkx2.1* | CTCATCGACATGATTCGGCG | TGCAACGGATCTGCGGG |
| *Nrp1* | AATGTCTTGTGAGAGCCCCAG | AGTGGCACAGGTGATGACTTC |

| | | |
|---|---|---|
| *Nrp2* | TTTGCCAGATGAGGGGTCAC | CAGCTTTTGCAGGTGAGGAT |
| *Npas1* | CCCCAGGATGGATGTAGTCG | GGCTTCGTGTTCGCTTTGAA |
| *Npy* | TTGTTCTGGGGGCGTTTTCT | TGGCCAGATACTACTCCGCT |
| *Nr2f2* | CCGTTGGTCAGGGCAAACT | GCGGAGGAACCTGAGCTACAC |
| *Nxph1* | TGGATTTGCTGCTCCCTGAC | GTTCTTCTCCTCCTGCAGCC |
| *Olig1* | TATAAGCCTGCGCTACGACG | CTCGCCCAGGTGTTTTGTTG |
| *Olig2* | AGATCATCGGGTTCTGGGGA | GCTTAGATCATCCCTGGGGC |
| *Prokr2* | GGGAGAACAGATACCATCCAGG | CCAAGTAGGCAAGCCTCAAC |
| *Reln* | CACGAGCTGCCAGGAATCGGAC | TGGCAACCCATCCTTCCACCTCT |
| *Rpp25* | CTCCTCTTGGACGTCGACAC | TCCACGACAACTTGGCTACC |
| *Slit1* | TACCTAGATGCTCCCAGCGA | TGCCCATTTTCCTCCTCACA |
| *Slit2* | CTCAGGCAGATTTGTGGGGA | AACGAGAGTTTGTCTGCAGTG |
| *Sox6* | GTGGATTTTGCTTAGCCGGG | GCCCGTGTCTACAGGGATG |
| *Sst* | GGACAGCATCTTCCTTGCCT | TGAAGGAGACGCTACCGAAG |
| *Tcf4* | GCCATTCGCTGTTGGTGATG | AAACCGAGCCAGGTGCATAA |

**Supplementary Table 2.** List of interneuron markers with their identity and reference source

| Gene Name | Identity 1 | Ref. | Identity 2 | Ref. | Identity 3 | Ref. | Identity 4 | Ref. |
|---|---|---|---|---|---|---|---|---|
| **Sept5** | Progenitor SST-E14.5 | Mi et al. | | | | | | |
| **1110001a16Rik** | GABAergic neuron development - SST | Mi et al. | | | | | | |
| **1110031i02Rik** | CGE-Progenitor | Mi et al. | | | | | | |
| **1500002o20Rik** | GABAergic neuron development - VIP | Mi et al. | | | | | | |
| **1500009L16Rik** | GABAergic neuron development - SST | Mayer et al. | | | | | | |
| **1700017b05Rik** | vMGE- progenitor | Mi et al. | | | | | | |
| **1810022k09Rik** | GABAergic neuron development - SST | Mi et al. | | | | | | |
| **2010007h12Rik** | GABAergic neuron development - VIP | Mi et al. | | | | | | |
| **2010204k13Rik** | GABAergic neuron development - VIP | Mi et al. | | | | | | |
| **2310028o11Rik** | GABAergic neuron development - SST | Mi et al. | | | | | | |
| **2310033p09Rik** | GABAergic neuron development - VIP | Mi et al. | | | | | | |
| **2900053a13Rik** | GABAergic neuron development - VIP | Mi et al. | | | | | | |
| **3110052m02Rik** | Progenitor SST-E14.5 | Mi et al. | | | | | | |
| **3110082i17Rik** | Progenitor SST-E14.5 | Mi et al. | | | | | | |
| **4921513d23Rik** | dMGE-CIN | Mi et al. | | | | | | |
| **4933415a04Rik** | vMGE-CIN-mature | Mi et al. | | | | | | |
| **5730577i03Rik** | CGE-Progenitor | Mi et al. | | | | | | |
| **5930434b04Rik** | Progenitor PV-E14.5 | Mi et al. | | | | | | |
| **7sk** | dMGE-CIN | Mi et al. | | | | | | |
| **9030425e11Rik** | Progenitor SST-E14.5 | Mi et al. | GABAergic neuron development - VIP | Mi et al. | | | | |
| **A730020e08Rik** | vMGE- progenitor | Mi et al. | | | | | | |
| **Aagab** | GABAergic neuron development - VIP | Mi et al. | | | | | | |

| | | | | |
|---|---|---|---|---|
| **Abcd4** | dMGE-progenitor | Mi et al. | | |
| **Abt1** | GABAergic neuron development - VIP | Mi et al. | | |
| **Acap3** | GABAergic neuron development - SST | Mi et al. | | |
| **Ackr3** | MGE-CIN | Chen et al. | | |
| **Acsl4** | dMGE-CIN | Mi et al. | | |
| **Acss1** | GABAergic neuron development - PV | Mi et al. | | |
| **Acvr2a** | Progenitor SST-E14.5 | Mi et al. | | |
| **Adarb2** | GABAergic neuron development - VIP | Mi et al. | | |
| **Adcy1** | GABAergic neuron development - VIP | Mi et al. | | |
| **Afap1** | Progenitor PV-E14.5 | Mi et al. | | |
| **Afg3l1 (Afg3l1p)** | Progenitor SST-E14.5 | Mi et al. | | |
| **Ahnak** | dMGE-progenitor | Mi et al. | | |
| **Amot** | GABAergic neuron development - SST | Mi et al. | | |
| **Amz2** | GABAergic neuron development - SST | Mi et al. | | |
| **Anks1b** | Progenitor SST-E14.5 | Mi et al. | | |
| **Anxa2** | Progenitor SST-E14.5 | Mi et al. | | |
| **Ap2a2** | Progenitor PV-E14.5 | Mi et al. | | |
| **Ap3b1** | GABAergic neuron development - SST | Mi et al. | | |
| **Aprt** | dMGE-progenitor | Mi et al. | | |
| **Arhgap33** | Progenitor PV-E14.5 | Mi et al. | | |
| **Armc10** | Progenitor SST-E14.5 | Mi et al. | | |
| **Armc8** | GABAergic neuron development - VIP | Mi et al. | | |
| **Armcx2** | GABAergic neuron development - VIP | Mi et al. | | |
| **Arx** | MGE-immature neurons | Chen et al. | Migration to Cortex (tangential) | Faux et al. |

| | | | | |
|---|---|---|---|---|
| **Arxes1** | GABAergic neuron development - VIP | Mi et al. | | |
| **Asap1** | CGE-derived migrating cells | Asgarian et al. | | |
| **Asb4** | GABAergic neuron development - PV | Mi et al. | vMGE-CIN | Mi et al. |
| **Aspm** | Progenitor | Mi et al. | | |
| **Atad2b** | dMGE-CIN | Mi et al. | | |
| **Atcay** | Markers previously not found in GABAergic neurons | Chen et al. | | |
| **Atg2b** | vMGE-CIN | Mi et al. | | |
| **Atp1a3** | LGE-immature neurons | Chen et al. | | |
| **Atp5e** | LGE-immature neurons | Chen et al. | | |
| **Atp5j** | GABAergic neuron development - SST | Mi et al. | | |
| **Atrip** | vMGE- progenitor | Mi et al. | | |
| **B3galnt1** | GABAergic neuron development - VIP | Mi et al. | | |
| **B3galt1** | GABAergic neuron development - VIP | Mi et al. | | |
| **Bbs9** | dMGE-progenitor | Mi et al. | | |
| **Bbx** | Progenitor PV-E14.5 | Mi et al. | | |
| **Bc057079** | vMGE- progenitor | Mi et al. | | |
| **Bcan** | vMGE - progenitor (PV+CINs) | Chen et al. | | |
| **Bcl11a** | GABAergic neuron development - SST | Mayer et al. | | |
| **Bcl7b** | Progenitor SST-E14.5 | Mi et al. | | |
| **Brcc3** | GABAergic neuron development - SST | Mi et al. | | |
| **Brwd1** | MGE-immature neurons | Chen et al. | | |
| **Btbd11** | CGE-derived migrating cells | Asgarian et al. | | |

| | | | | |
|---|---|---|---|---|
| **C030044b11Rik** | CGE-Progenitor | Mi et al. | | |
| **Cacna2d3** | GABAergic neuron development - SST | Mayer et al. | | |
| **Cacnb3** | Progenitor PV-E14.5 | Mi et al. | | |
| **Cacng3** | MGE-derived IN | Mayer et al. | | |
| **Cadm2** | CGE-derived migrating cells | Asgarian et al. | | |
| **Calb1** | GABAergic neuron development - SST | Mi et al. | | |
| **Cbfa2t3** | MGE-CIN | Asgarian et al. | MGE-derived migrating cells | Asgarian et al. |
| **Ccdc23** | GABAergic neuron development - SST | Mi et al. | | |
| **Ccdc23 (Svbp)** | Progenitor SST-E14.5 | Mi et al. | | |
| **Ccdc90b** | Progenitor SST-E14.5 | Mi et al. | | |
| **Ccnd2** | Progenitor | Mi et al. | GABAergic neuron development - PV | Mi et al. |
| **Ccne2** | Progenitor | Mi et al. | | |
| **Ccnl1** | MGE-immature neurons | Chen et al. | | |
| **Cdc6** | Progenitor | Mi et al. | | |
| **Cdca7** | GABAergic neuron development - PV | Mi et al. | | |
| **Cdk14** | GABAergic neuron development - SST | Mi et al. | | |
| **Cdkn1c** | LGE-immature neurons | Chen et al. | | |
| **Cds2** | Progenitor SST-E14.5 | Mi et al. | | |
| **Cebpz** | vMGE-CIN | Mi et al. | | |
| **Celsr1** | Progenitor SST-E14.5 | Mi et al. | | |
| **Cenpe** | Progenitor | Mi et al. | | |
| **Cep120** | MGE-immature neurons | Chen et al. | | |
| **Cfsa** | GABAergic neuron development - PV | Mi et al. | | |
| **Clspn** | Progenitor | Mi et al. | | |

| | | | | | | |
|---|---|---|---|---|---|---|
| **Clstn1** | GABAergic neuron development - VIP | Mi et al. | | | | |
| **Cmas** | GABAergic neuron development - SST | Mi et al. | | | | |
| **Cmtm6** | CGE-Progenitor | Mi et al. | | | | |
| **Cno** | Progenitor SST-E14.5 | Mi et al. | | | | |
| **Cog6** | GABAergic neuron development - VIP | Mi et al. | | | | |
| **Col22a1** | Progenitor SST-E14.5 | Mi et al. | | | | |
| **Col5a2** | vMGE-CIN | Mi et al. | | | | |
| **Cops4** | GABAergic neuron development - SST | Mi et al. | | | | |
| **Coup-TFI** | CGE specification | Faux et al. | | | | |
| **Cox19** | GABAergic neuron development - SST | Mi et al. | | | | |
| **Cox5a** | MGE-derived migrating cells | Asgarian et al. | | | | |
| **Cox6c** | LGE-immature neurons | Chen et al. | | | | |
| **Cox7a2l** | GABAergic neuron development - SST | Mi et al. | | | | |
| **Cplx1** | PV-CINs | Chen et al. | | | | |
| **Cpsf6** | GABAergic neuron development - SST | Mi et al. | | | | |
| **Crabp1** | LGE-immature neurons | Chen et al. | GABAergic neuron development - VIP | Mi et al. | | |
| **Crabp2** | MGE-immature neurons | Chen et al. | GABAergic neuron development - PV | Mi et al. | vMGE-CIN | Mi et al. |
| **Crebl2** | vMGE-CIN | Mi et al. | | | | |
| **Cuedc1** | Progenitor SST-E14.5 | Mi et al. | | | | |
| **Cux1** | immature neurons | Faux et al | | | | |
| **Cux2** | immature neurons | Faux et al | | | | |
| **Cwc22** | dMGE-CIN | Mi et al. | | | | |
| **Cxcl14** | CGE-derived migrating cells | Asgarian et al. | GABAergic neuron development - VIP | Mi et al. | | |
| **D16h22s680e** | Progenitor SST-E14.5 | Mi et al. | | | | |

| | | | | | | |
|---|---|---|---|---|---|---|
| **D17h6s53e** | dMGE-CIN | Mi et al. | | | | |
| **Dcbld1** | Progenitor SST-E14.5 | Mi et al. | | | | |
| **Dcc** | MGE-immature neurons | Chen et al. | | | | |
| **Dchs1** | GABAergic neuron development - SST | Mi et al. | | | | |
| **Dclk2** | Progenitor PV-E14.5 | Mi et al. | | | | |
| **Dclre1b** | GABAergic neuron development - PV | Mi et al. | dMGE-CIN | Mi et al. | | |
| **Dcx** | Maturing/postmitotic neurons | Chen et al. | Migration to Cortex (tangential) | Faux et al. | | |
| **Ddah1** | Progenitor | Mi et al. | | | | |
| **Ddb1** | GABAergic neuron development - PV | Mi et al. | | | | |
| **Ddhd1** | GABAergic neuron development - VIP | Mi et al. | | | | |
| **Ddt** | vMGE- progenitor | Mi et al. | | | | |
| **Dennd4b** | vMGE- progenitor | Mi et al. | | | | |
| **Dhx8** | GABAergic neuron development - PV | Mi et al. | | | | |
| **Dleu2** | GABAergic neuron development - VIP | Mi et al. | | | | |
| **Dlg2** | CGE-derived migrating cells | Asgarian et al. | | | | |
| **Dlx1** | MGE-immature neurons | Chen et al., Wonders et al. | Migration to Cortex (tangential) | Wonders et al. | | |
| **Dlx2** | MGE-immature neurons | Chen et al., Wonders et al. | Migration to Cortex (tangential) | Wonders et al. | | |
| **Dlx5** | MGE-immature neurons | Wonders et al. | GABAergic neuron development | Chen et al. | Migration to Cortex (tangential) | Faux et al. |
| **Dlx6** | MGE-immature neurons | Wonders et al. | Migration to Cortex (tangential) | Wonders et al. | Cortical migration (radial) | Wonders et al. |
| **Dlx6os1** | Migration to Cortex (tangential) | Wonders et al. | Cortical migration (radial) | Wonders et al | | |
| **Dmxl2** | GABAergic neuron development - VIP | Mi et al. | | | | |
| **Dnahc7b** | dMGE-CIN | Mi et al. | | | | |

| | | | | | | |
|---|---|---|---|---|---|---|
| **Dnajb9** | Progenitor SST-E14.5 | Mi et al. | | | | |
| **Dnd1** | GABAergic neuron development - VIP | Mi et al. | | | | |
| **Dock6** | vMGE-CIN | Mi et al. | | | | |
| **Dpysl2** | MGE-immature neurons | Chen et al. | | | | |
| **Dtd1** | GABAergic neuron development - VIP | Mi et al. | | | | |
| **Dusp7** | dMGE-CIN | Mi et al. | | | | |
| **Dyrk3** | CGE-Progenitor | Mi et al. | | | | |
| **Ebf1** | CGE-Progenitor | Mi et al. | | | | |
| **Ece1** | vMGE-CIN | Mi et al. | | | | |
| **Edem2** | GABAergic neuron development - SST | Mi et al. | | | | |
| **Ednrb** | Progenitor | Mi et al. | | | | |
| **Eif2c1** | Progenitor PV-E14.5 | Mi et al. | | | | |
| **Eif5** | GABAergic neuron development - SST | Mi et al. | | | | |
| **Eif5a2** | vMGE- progenitor | Mi et al. | | | | |
| **Eme1** | GABAergic neuron development - PV | Mi et al. | | | | |
| **Epb4.1l1** | GABAergic neuron development - SST | Mi et al. | | | | |
| **Epb4.1l4a** | GABAergic neuron development - VIP | Mi et al. | | | | |
| **Epdr1** | CGE-Progenitor | Mi et al. | | | | |
| **Epha5** | GABAergic neuron development - SST | Mi et al. | | | | |
| **Ephb1** | LGE-immature neurons | Chen et al. | | | | |
| **Ephb3** | MGE-SIN | Chen et al. | | | | |
| **Erbb4** | MGE-immature neurons | Chen et al. | GABAergic neuron development - SST | Mi et al. | MGE-SIN | Chen et al. |
| **Etv1** | vMGE - progenitor (PV+CINs) | Chen et al. | | | | |
| **Evi2b** | vMGE-CIN | Mi et al. | | | | |

| | | | | |
|---|---|---|---|---|
| **Exosc5** | GABAergic neuron development - PV | Mi et al. | | |
| **Ezr** | GABAergic neuron development - PV | Mi et al. | | |
| **Fabp7** | Progenitor | Mi et al. | Progenitor SST-E14.5 | Mi et al. |
| **Faim2** | Progenitor | Mi et al. | | |
| **Fam117b** | GABAergic neuron development - VIP | Mi et al. | | |
| **Fapb7** | Progenitor | Mi et al. | | |
| **Fat4** | dMGE-CIN | Mi et al. | | |
| **Fbrsl1** | Progenitor PV-E14.5 | Mi et al. | | |
| **Fbxo9** | GABAergic neuron development - VIP | Mi et al. | | |
| **Fign** | MGE-immature neurons | Chen et al. | | |
| **Flrt2** | GABAergic neuron development - VIP | Mi et al. | | |
| **Fnta** | Progenitor SST-E14.5 | Mi et al. | | |
| **Foxg1** | MGE-immature neurons | Chen et al. | | |
| **FoxJ1** | vMGE - progenitor (PV+CINs) | Chen et al. | | |
| **Foxn3** | MGE-immature neurons | Chen et al. | | |
| **Foxp2** | Progenitor SST-E14.5 | Mi et al. | GABAergic neuron development - VIP | Mi et al. |
| **Foxred1** | GABAergic neuron development - VIP | Mi et al. | | |
| **Frmpd1** | Progenitor | Mi et al. | | |
| **Fxyd6** | GABAergic neuron development - SST | Mayer et al. | | |
| **Gabrb3** | GABAergic neuron development - VIP | Mi et al. | | |
| **Gabrg2** | CGE-Progenitor | Mi et al. | | |
| **Gad1** | GABAergic neuron development | Chen et al. | | |

| | | | | | | |
|---|---|---|---|---|---|---|
| **Gad2** | GABAergic neuron development | Chen et al. | | | | |
| **Gap43** | LGE-immature neurons | Chen et al. | Maturing/postmitotic neurons | Chen et al. | MGE-CIN | Chen et al. |
| **Gem** | dMGE-progenitor | Mi et al. | | | | |
| **Gfap** | Progenitor | Mi et al. | | | | |
| **Gli1** | dMGE - progenior (STT+CINs) | Chen et al. | | | | |
| **Gli2** | dMGE - progenior (STT+CINs) | Chen et al. | | | | |
| **Glipr2** | GABAergic neuron development - SST | Mi et al. | | | | |
| **Glrx3** | GABAergic neuron development - VIP | Mi et al. | | | | |
| **Glrx5** | CGE-Progenitor | Mi et al. | Progenitor SST-E14.5 | Mi et al. | | |
| **Gm10277** | dMGE-progenitor | Mi et al. | dMGE-CIN | Mi et al. | | |
| **Gm10561** | dMGE-CIN | Mi et al. | | | | |
| **Gm10837** | dMGE-progenitor | Mi et al. | | | | |
| **Gm11934** | GABAergic neuron development - SST | Mi et al. | | | | |
| **Gm12355** | GABAergic neuron development - SST | Mi et al. | | | | |
| **Gm14964** | Progenitor PV-E14.5 | Mi et al. | | | | |
| **Gm15501** | Progenitor PV-E14.5 | Mi et al. | | | | |
| **Gm16076** | dMGE-CIN | Mi et al. | | | | |
| **Gm16869** | Progenitor PV-E14.5 | Mi et al. | | | | |
| **Gm17284** | vMGE-CIN | Mi et al. | | | | |
| **Gm17608** | dMGE-CIN | Mi et al. | | | | |
| **Gm17635** | vMGE- progenitor | Mi et al. | | | | |
| **Gm4540** | GABAergic neuron development - SST | Mi et al. | | | | |
| **Gm6531** | GABAergic neuron development - SST | Mi et al. | | | | |
| **Gng3** | Maturing/postmitotic neurons | Chen et al. | GABAergic neuron development - SST | Mi et al. | | |
| **Gpr98** | Progenitor | Mi et al. | | | | |

| | | | | | |
|---|---|---|---|---|---|
| **Gprasp1** | GABAergic neuron development - VIP | Mi et al. | | | |
| **Gria2** | LGE-immature neurons | Chen et al. | Maturing/postmitotic neurons | Chen et al. | Mi et al. |
| **Gria3** | dMGE-progenitor | Mi et al. | | | |
| **Gria4** | GABAergic neuron development - SST | Mi et al. | | | |
| **Grtp1** | GABAergic neuron development - PV | Mi et al. | | | |
| **Gsk3b** | MGE-immature neurons | Chen et al. | | | |
| **Gt(Rosa)26sor** | Progenitor SST-E14.5 | Mi et al. | | | |
| **Gtpbp4** | GABAergic neuron development - SST | Mi et al. | | | |
| **H2-ke2** | GABAergic neuron development - SST | Mi et al. | | | |
| **Hace1** | Progenitor SST-E14.5 | Mi et al. | | | |
| **Hat1** | Progenitor | Mi et al. | | | |
| **Hbp1** | MGE-immature neurons | Chen et al. | | | |
| **Heatr5a** | vMGE- progenitor | Mi et al. | | | |
| **Hells** | Progenitor | Mi et al. | | | |
| **Helt** | Progenitor SST-E14.5 | Mi et al. | | | |
| **Hes1** | Progenitor | Mi et al. | | | |
| **Hes5** | Progenitor | Mi et al. | | | |
| **Hhip** | dMGE - progenior (STT+CINs) | Chen et al. | | | |
| **Hint1** | GABAergic neuron development - SST | Mi et al. | | | |
| **Hint2** | GABAergic neuron development - SST | Mi et al. | | | |
| **Hist1h1a** | dMGE-CIN | Mi et al. | | | |
| **Hist1h2ak** | GABAergic neuron development - VIP | Mi et al. | | | |
| **Htr3a** | CGE-derived migrating cells | Asgarian et al. | | | |

| | | | | | | |
|---|---|---|---|---|---|---|
| **Id2** | GABAergic neuron development - SST | Mi et al. | | | | |
| **Ids** | GABAergic neuron development - VIP | Mi et al. | | | | |
| **Ift20** | Progenitor PV-E14.5 | Mi et al. | | | | |
| **Igfbpl1** | Progenitor SST-E14.5 | Mi et al. | | | | |
| **Ilf3** | MGE-immature neurons | Chen et al. | | | | |
| **Ina** | Progenitor SST-E14.5 | Mi et al. | GABAergic neuron development - VIP | Mi et al. | | |
| **Ionrf1** | vMGE- progenitor | Mi et al. | | | | |
| **Ipo11** | Progenitor SST-E14.5 | Mi et al. | | | | |
| **Iqce** | dMGE-progenitor | Mi et al. | | | | |
| **Iscu** | Progenitor SST-E14.5 | Mi et al. | | | | |
| **Isl1** | LGE-immature neurons | Chen et al. | | | | |
| **Itga7** | Progenitor PV-E14.5 | Mi et al. | | | | |
| **Kctd2** | CGE-Progenitor | Mi et al. | | | | |
| **Kdm3b** | GABAergic neuron development - VIP | Mi et al. | | | | |
| **Kdm5a** | MGE-immature neurons | Chen et al. | | | | |
| **Kif20a** | GABAergic neuron development - PV | Mi et al. | | | | |
| **Kif21b** | Progenitor PV-E14.5 | Mi et al. | | | | |
| **Kif22** | Progenitor | Mi et al. | | | | |
| **Kif7** | dMGE-CIN | Mi et al. | | | | |
| **L1cam** | MGE-immature neurons | Chen et al. | Maturing/postmitotic neurons | Chen et al. | | |
| **Ldb1** | Progenitor PV-E14.5 | Mi et al. | | | | |
| **Lfng** | Progenitor SST-E14.5 | Mi et al. | | | | |
| **Lhx2** | Progenitor | Mi et al. | | | | |
| **Lhx6** | MGE-immature neurons | Chen et al., Wonders et al. | Migration to Cortex | Wonders et al. | GABAergic neuron development | Chen et al. |
| **Lhx8** | vMGE- progenitor | Mi et al. | GABAergic neuron development | Chen et al., Mi et al. | | |

| | | | | |
|---|---|---|---|---|
| **Lrpprc** | GABAergic neuron development - VIP | Mi et al. | | |
| **Lrrc57** | GABAergic neuron development - PV | Mi et al. | | |
| **Ly6h** | GABAergic neuron development - SST | Mayer et al. | | |
| **Lyar** | GABAergic neuron development - VIP | Mi et al. | | |
| **Maf** | MGE-immature neurons | Chen et al. | GABAergic neuron development - SST | Mi et al. |
| **Mafb** | MGE-immature neurons | Chen et al. | | |
| **Map2** | Maturing/postmitotic neurons | Chen et al. | | |
| **Map3k9** | CGE-Progenitor | Mi et al. | | |
| **Mapt** | LGE-immature neurons | Chen et al. | Maturing/postmitotic neurons | Chen et al., Mi et al. |
| **Mash1** | immature neurons | Faux et al. | | |
| **Maz** | GABAergic neuron development - VIP | Mi et al. | | |
| **Mbtd1** | Progenitor PV-E14.5 | Mi et al. | | |
| **Mcm3** | Progenitor | Mi et al. | | |
| **Mcm4** | Progenitor | Mi et al. | | |
| **Mcm6** | Progenitor | Mi et al. | | |
| **Mdm2** | MGE-immature neurons | Chen et al. | | |
| **Med12** | dMGE-progenitor | Mi et al. | | |
| **Mef2c** | GABAergic neuron development - PV | Mayer et al. | | |
| **Meis2** | LGE-immature neurons | Chen et al. | GABAergic neuron development - VIP | Mi et al. |
| **Mex3c** | vMGE-CIN | Mi et al. | | |
| **Mfng** | GABAergic neuron development - PV | Mi et al. | | |
| **Mif** | LGE-immature neurons | Chen et al. | | |
| **Mki67** | Progenitor | Mi et al. | | |

| | | | | |
|---|---|---|---|---|
| **Mknk2** | dMGE-progenitor | Mi et al. | | |
| **Mllt11** | Markers previously not found in GABAergic neurons | Chen et al. | | |
| **Mllt3** | Markers previously not found in GABAergic neurons | Chen et al. | | |
| **Mllt4 (Afdn)** | Progenitor PV-E14.5 | Mi et al. | | |
| **Mns1** | GABAergic neuron development - PV | Mi et al. | | |
| **Mpp2** | dMGE-CIN | Mi et al. | | |
| **Mrrf** | GABAergic neuron development - PV | Mi et al. | | |
| **Msh2** | GABAergic neuron development - VIP | Mi et al. | | |
| **Msi1** | MGE-immature neurons | Chen et al. | | |
| **Msi2** | MGE-immature neurons | Chen et al. | | |
| **Mt2** | GABAergic neuron development - PV | Mi et al. | vMGE-CIN | Mi et al. |
| **Mt3** | Progenitor SST-E14.5 | Mi et al. | | |
| **Mthfr** | GABAergic neuron development - PV | Mi et al. | | |
| **Myeov2** | Progenitor PV-E14.5 | Mi et al. | | |
| **Myh9** | Progenitor PV-E14.5 | Mi et al. | | |
| **Myo10** | Progenitor | Mi et al. | | |
| **Myt1** | Markers previously not found in GABAergic neurons | Chen et al. | | |
| **Nasp** | Progenitor | Mi et al. | | |
| **Nat15** | GABAergic neuron development - SST | Mi et al. | | |
| **Nav2** | Progenitor PV-E14.5 | Mi et al. | | |
| **Nbr1** | GABAergic neuron development - PV | Mi et al. | | |
| **Nckap5** | Progenitor SST-E14.5 | Mi et al. | | |

| | | | | |
|---|---|---|---|---|
| **Ndc60** | GABAergic neuron development - SST | Mi et al. | | |
| **Nde1** | Progenitor | Mi et al. | | |
| **Ndfip1** | Progenitor SST-E14.5 | Mi et al. | | |
| **Ndn** | LGE-immature neurons | Chen et al. | | |
| **Ndst3** | CGE-Progenitor | Mi et al. | | |
| **Ndufb2** | Progenitor PV-E14.5 | Mi et al. | | |
| **Ndufb4** | GABAergic neuron development - SST | Mi et al. | | |
| **Ndufb8** | GABAergic neuron development - SST | Mi et al. | | |
| **Ndufs2** | GABAergic neuron development - SST | Mi et al. | | |
| **Nenf** | GABAergic neuron development - SST | Mayer et al. | | |
| **Neto1** | GABAergic neuron development - SST | Mayer et al. | | |
| **Nfia** | MGE-immature neurons | Chen et al. | | |
| **Nfix** | Progenitor SST-E14.5 | Mi et al. | | |
| **Ngfrap1** | GABAergic neuron development - SST | Mi et al. | | |
| **Nkiras2** | GABAergic neuron development - VIP | Mi et al. | | |
| **Nkx2-1** | GABAergic neuron development - PV | Mi et al. | | |
| **Nkx6-2** | dMGE - progenior (STT-CINs) | Chen et al. | MGE-Interneurons (specification | Faux et al. |
| **Nlng1** | GABAergic neuron development - VIP | Mi et al. | | |
| **Nmd3** | GABAergic neuron development - SST | Mi et al. | | |
| **Nop10** | GABAergic neuron development - SST | Mi et al. | | |
| **Notch1** | Progenitor | Mi et al. | | |
| **Notch2** | Progenitor | Mi et al. | | |

| | | | | |
|---|---|---|---|---|
| **Notch3** | Progenitor | Mi et al. | | |
| **Npas1** | GABAergic neuron development - VIP | Mi et al. | | |
| **Npy** | STT-CINs | Chen et al. | GABAergic neuron development - SST | Mi et al. |
| **Nr2f1** | dMGE - progenior (STT-CINs) | Chen et al. | | |
| **Nr2f2** | GABAergic neuron development - SST | Mi et al. | | |
| **Nrg1** | LGE-immature neurons | Chen et al. | | |
| **Nrp2** | GABAergic neuron development - VIP | Mi et al. | | |
| **Nrxn1** | LGE-immature neurons | Chen et al. | | |
| **Nrxn3** | Progenitor PV-E14.5 | Mi et al. | Maturing/postmitotic neurons | Chen et al. |
| **Ntrk3** | LGE-immature neurons | Chen et al. | | |
| **Nxph1** | MGE-CIN | Chen et al. | GABAergic neuron development - SST | Mi et al. |
| **Nxph2** | MGE-immature neurons | Mayer et al. | | |
| **Olig1** | Progenitor | Mi et al. | | |
| **Pabpc4** | Progenitor SST-E14.5 | Mi et al. | | |
| **Pan3** | GABAergic neuron development - PV | Mi et al. | | |
| **Papolg** | GABAergic neuron development - VIP | Mi et al. | | |
| **Parp6** | Progenitor SST-E14.5 | Mi et al. | | |
| **Parvb** | MGE-CIN | Asgarian et al. | | |
| **Pask** | dMGE-CIN | Mi et al. | | |
| **Pax6** | GABAergic neuron development - VIP | Mi et al. | | |
| **Pbx3** | Progenitor SST-E14.5 | Mi et al. | | |
| **Pcnxl3** | GABAergic neuron development - PV | Mi et al. | | |

| | | |
|---|---|---|
| **Pcyt1a** | Progenitor PV-E14.5 | Mi et al. |
| **Pdcd7** | dMGE-progenitor | Mi et al. |
| **Pde1a** | STT-CINs | Chen et al. |
| **Pde4b** | GABAergic neuron development - VIP | Mi et al. |
| **Pdlim5** | vMGE- progenitor | Mi et al. |
| **Pdpn** | Progenitor | Mi et al. |
| **Pfas** | Progenitor | Mi et al. |
| **Pfdn4** | GABAergic neuron development - VIP | Mi et al. |
| **Pfkm** | GABAergic neuron development - PV | Mi et al. |
| **Pgap1** | Progenitor SST-E14.5 | Mi et al. |
| **Phax** | Progenitor PV-E14.5 | Mi et al. |
| **Phf2** | dMGE-progenitor | Mi et al. |
| **Phf6** | GABAergic neuron development - SST | Mi et al. |
| **Phospho2** | Progenitor SST-E14.5 | Mi et al. |
| **Phyh** | GABAergic neuron development - PV | Mi et al. |
| **Pip5kl1** | Progenitor | Mi et al. |
| **Pisd-ps2** | GABAergic neuron development - SST | Mi et al. |
| **Pitpnc1** | GABAergic neuron development - SST | Mi et al. |
| **Plcl2** | CGE-Progenitor | Mi et al. |
| **Plcxd3** | GABAergic neuron development - PV | Mayer et al. |
| **Plin3** | dMGE-CIN | Mi et al. |
| **Plxna1** | Maturing/postmitotic neurons | Chen et al. |
| **Polr2a** | vMGE-CIN | Mi et al. |
| **Pou3f2** | vMGE-CIN | Mi et al. |
| **Pou3f3** | MGE-immature neurons | Chen et al. |

| | | |
|---|---|---|
| **Pp1r1a** | GABAergic neuron development - VIP | Mi et al. |
| **Ppp1r15a** | GABAergic neuron development - PV | Mi et al. |
| **Ppp1r15b** | GABAergic neuron development - VIP | Mi et al. |
| **Ppp2r2b** | Progenitor SST-E14.5 | Mi et al. |
| **Ppp3r1** | Progenitor PV-E14.5 | Mi et al. |
| **Ppp4r1l-ps** | GABAergic neuron development - PV | Mi et al. |
| **Pqlc2** | GABAergic neuron development - VIP | Mi et al. |
| **Prkaca** | Progenitor SST-E14.5 | Mi et al. |
| **Prmt1** | Progenitor PV-E14.5 | Mi et al. |
| **Prox1** | CGE Progenitor | Mayer et al. |
| **Psmb5** | Progenitor PV-E14.5 | Mi et al. |
| **Psmb6** | GABAergic neuron development - SST | Mi et al. |
| **Psmc4** | Progenitor | Mi et al. |
| **Ptch1** | vMGE- progenitor | Mi et al. |
| **Ptprd** | GABAergic neuron development - SST | Mi et al. |
| **Ptpru** | GABAergic neuron development - VIP | Mi et al. |
| **Pvalb** | PV-CINs | Chen et al. |
| **Rab3b** | GABAergic neuron development - SST | Mayer et al. |
| **Rab5a** | GABAergic neuron development - PV | Mi et al. |
| **Rad17** | GABAergic neuron development - SST | Mi et al. |
| **Rad54t2** | GABAergic neuron development - SST | Mi et al. |
| **Ralgps2** | GABAergic neuron development - SST | Mi et al. |
| **Ranbp1** | GABAergic neuron development - SST | Mi et al. |

| | | | | | | | | |
|---|---|---|---|---|---|---|---|---|
| **Rap1b** | CGE-Progenitor | Mi et al. | | | | | | |
| **Rbm15** | GABAergic neuron development - SST | Mi et al. | | | | | | |
| **Reln** | LGE-immature neurons | Chen et al. | STT-CINs | Chen et al. | | | | |
| **Rftn2** | GABAergic neuron development - PV | Mi et al. | | | | | | |
| **Rfwd3** | GABAergic neuron development - SST | Mi et al. | | | | | | |
| **Rgs17** | GABAergic neuron development - SST | Mayer et al. | | | | | | |
| **Rhpn1** | Progenitor SST-E14.5 | Mi et al. | | | | | | |
| **Rnasel** | Progenitor PV-E14.5 | Mi et al. | | | | | | |
| **Rnf214** | GABAergic neuron development - SST | Mi et al. | GABAergic neuron development - VIP | Mi et al. | | | | |
| **Rnf4** | MGE-immature neurons | Chen et al. | | | | | | |
| **Rpl41** | LGE-immature neurons | Chen et al. | | | | | | |
| **Rpp25** | MGE-derived migrating cells | Asgarian et al. | | | | | | |
| **Rrm2** | Progenitor | Mi et al. | GABAergic neuron development - PV | Mi et al. | | | | |
| **Rtn1** | GABAergic neuron development - SST | Mayer et al. | | | | | | |
| **Rtn4** | GABAergic neuron development - SST | Mi et al. | | | | | | |
| **Runx1t1** | MGE-immature neurons | Chen et al. | Progenitor PV-E14.5 | Mi et al. | MGE-derived Cortical SST | Asgarian et al. | Markers previously not found in GABAergic neurons | Chen et al. |
| **Sall3** | Progenitor | Mi et al. | | | | | | |
| **Satb1** | MGE-immature neurons | Chen et al. | | | | | | |
| **Satb2** | Progenitor PV-E14.5 | Mi et al. | | | | | | |
| **Scamp1** | GABAergic neuron development - SST | Mi et al. | | | | | | |
| **Scamp5** | GABAergic neuron development - SST | Mi et al. | | | | | | |

| | | | | |
|---|---|---|---|---|
| **Scn3a** | LGE-immature neurons | Chen et al. | Maturing/postmitotic neurons | Chen et al. |
| **Sema3c** | dMGE-progenitor | Mi et al. | | |
| **Sema5b** | GABAergic neuron development - SST | Mi et al. | | |
| **Serbp1** | GABAergic neuron development - SST | Mi et al. | | |
| **Serinc3** | Progenitor PV-E14.5 | Mi et al. | | |
| **Sesn3** | vMGE- progenitor | Mi et al. | | |
| **Setbp1** | MGE-immature neurons | Chen et al. | | |
| **Ska1** | Progenitor | Mi et al. | | |
| **Slc1a3** | Progenitor | Mi et al. | Progenitor SST-E14.5 | Mi et al. |
| **Slc25a36** | GABAergic neuron development - VIP | Mi et al. | | |
| **Slc48a1** | GABAergic neuron development - VIP | Mi et al. | | |
| **Slc8a1** | GABAergic neuron development - VIP | Mi et al. | | |
| **Slc9a9** | dMGE-progenitor | Mi et al. | | |
| **Slx1b** | vMGE-CIN | Mi et al. | | |
| **Slx4** | dMGE-progenitor | Mi et al. | | |
| **Smarcd3** | Progenitor SST-E14.5 | Mi et al. | | |
| **Smn1** | Progenitor PV-E14.5 | Mi et al. | | |
| **Smpd2** | GABAergic neuron development - PV | Mi et al. | | |
| **Snap25** | LGE-immature neurons | Chen et al. | | |
| **Snap47** | LGE-immature neurons | Chen et al. | MGE-CIN | Chen et al. |
| **Sncalp** | Progenitor SST-E14.5 | Mi et al. | | |
| **Sntg1** | GABAergic neuron development - VIP | Mi et al. | | |
| **Sobp** | vMGE- progenitor | Mi et al. | | |
| **Socs2** | GABAergic neuron development - VIP | Mi et al. | | |

| | | | | | | |
|---|---|---|---|---|---|---|
| **Sox11** | MGE-immature neurons | Chen et al. | Progenitor PV-E14.5 | Mi et al. | | |
| **Sox2** | Progenitor | Mi et al. | MGE-CIN | Chen et al. | | |
| **Sox4** | MGE-immature neurons | Chen et al. | | | | |
| **Sox5** | PV-CINs | Chen et al. | | | | |
| **Sox6** | MGE-immature neurons | Chen et al. | | | | |
| **Sp140** | dMGE-progenitor | Mi et al. | | | | |
| **Sp8** | GABAergic neuron development - VIP | Mi et al. | | | | |
| **Sp9** | MGE-immature neurons | Chen et al. | | | | |
| **Sparc** | Progenitor | Mi et al. | Progenitor SST-E14.5 | Mi et al. | | |
| **Spink10** | GABAergic neuron development - VIP | Mi et al. | | | | |
| **Spock3** | GABAergic neuron development - VIP | Mi et al. | | | | |
| **Srebf2** | GABAergic neuron development - SST | Mi et al. | | | | |
| **Srsf4** | Progenitor SST-E14.5 | Mi et al. | | | | |
| **Ssb** | GABAergic neuron development - SST | Mi et al. | | | | |
| **Sst** | CGE-Progenitor | Mi et al. | GABAergic neuron development - SST | Mi et al. | STT-CINs | Chen et al. |
| **St6galnac6** | Progenitor SST-E14.5 | Mi et al. | | | | |
| **Stard5** | vMGE- progenitor | Mi et al. | | | | |
| **Stk33** | GABAergic neuron development - PV | Mi et al. | | | | |
| **Stmn2** | LGE-immature neurons | Chen et al. | Maturing/postmitotic neurons | Chen et al. | | |
| **Stmn3** | Maturing/postmitotic neurons | Chen et al. | | | | |
| **Str3a** | GABAergic neuron development - VIP | Mi et al. | | | | |
| **Stxbp1** | CGE-Progenitor | Mi et al. | | | | |

| | | | | |
|---|---|---|---|---|
| **Stxbp4** | GABAergic neuron development - VIP | Mi et al. | | |
| **Sulf1** | vMGE - progenitor (PV-CINs) | Chen et al. | | |
| **Sulf2** | vMGE - progenitor (PV-CINs) | Chen et al. | | |
| **Suv39h1** | Progenitor PV-E14.5 | Mi et al. | | |
| **Syt1** | GABAergic neuron development - SST | Mi et al. | | |
| **Syt2** | PV-CINs | Chen et al. | | |
| **Syt5** | CGE-Progenitor | Mi et al. | | |
| **Syt7** | GABAergic neuron development - VIP | Mi et al. | | |
| **Tacc3** | Progenitor | Mi et al. | | |
| **Tada1** | GABAergic neuron development - SST | Mi et al. | | |
| **Taf5l** | GABAergic neuron development - SST | Mi et al. | | |
| **Tbcc** | Progenitor SST-E14.5 | Mi et al. | | |
| **Tcf4** | MGE-immature neurons | Chen et al. | | |
| **Tctn2** | Progenitor SST-E14.5 | Mi et al. | | |
| **Thap1** | vMGE- progenitor | Mi et al. | | |
| **Tle4** | LGE-immature neurons | Chen et al. | | |
| **Tmeff2** | GABAergic neuron development - SST | Mayer et al. | | |
| **Tmem101** | GABAergic neuron development - VIP | Mi et al. | | |
| **Tmem130** | LGE-immature neurons | Chen et al. | Maturing/postmitotic neurons | Chen et al. |
| **Tmem165** | GABAergic neuron development - VIP | Mi et al. | | |
| **Tmem167b** | Progenitor PV-E14.5 | Mi et al. | | |
| **Tmem50a** | GABAergic neuron development - SST | Mi et al. | | |

| | | | | |
|---|---|---|---|---|
| **Tomm20** | GABAergic neuron development - SST | Mi et al. | | |
| **Tomm40** | Progenitor SST-E14.5 | Mi et al. | | |
| **Top2a** | Progenitor | Mi et al. | | |
| **Tox3** | Progenitor PV-E14.5 | Mi et al. | | |
| **Tpd52** | GABAergic neuron development - VIP | Mi et al. | | |
| **Tpr** | GABAergic neuron development - VIP | Mi et al. | | |
| **Tpst1** | GABAergic neuron development - VIP | Mi et al. | | |
| **Traf2** | GABAergic neuron development - PV | Mi et al. | | |
| **Trappc2l** | GABAergic neuron development - VIP | Mi et al. | | |
| **Trappc6a** | GABAergic neuron development - VIP | Mi et al. | | |
| **Trim11** | GABAergic neuron development - VIP | Mi et al. | | |
| **Trim13** | Progenitor SST-E14.5 | Mi et al. | | |
| **Trim46** | Progenitor PV-E14.5 | Mi et al. | | |
| **Tspan7** | GABAergic neuron development - SST | Mayer et al. | | |
| **Ttr** | vMGE- progenitor | Mi et al. | | |
| **Ttyh1** | Progenitor SST-E14.5 | Mi et al. | | |
| **Tubb2a** | GABAergic neuron development - SST | Mi et al. | | |
| **Tubb3** | Maturing/postmitotic neurons | Chen et al. | | |
| **U2af1** | GABAergic neuron development - SST | Mi et al. | | |
| **U4atac** | Progenitor PV-E14.5 | Mi et al. | | |
| **Uba52** | dMGE-CIN | Mi et al. | | |
| **Ube2ql1** | Progenitor SST-E14.5 | Mi et al. | GABAergic neuron development - SST | Mi et al. |
| **Ube2s** | CGE-Progenitor | Mi et al. | | |

| | | | | |
|---|---|---|---|---|
| **Ube3c** | GABAergic neuron development - SST | Mi et al. | | |
| **Uchl4** | GABAergic neuron development - PV | Mi et al. | | |
| **Uqcc** | GABAergic neuron development - PV | Mi et al. | | |
| **Usp24** | GABAergic neuron development - VIP | Mi et al. | | |
| **Vim** | Progenitor | Mi et al. | | |
| **Vldlr** | CGE-Progenitor | Mi et al. | Progenitor SST-E14.5 | Mi et al. |
| **Vti1b** | Progenitor SST-E14.5 | Mi et al. | | |
| **Wdr19** | vMGE- progenitor | Mi et al. | | |
| **Wdr91** | vMGE- progenitor | Mi et al. | | |
| **Whsc1l1** | GABAergic neuron development - VIP | Mi et al. | | |
| **Zc3h12c** | GABAergic neuron development - PV | Mi et al. | | |
| **Zc3h14** | GABAergic neuron development - VIP | Mi et al. | | |
| **Zdhhc2** | GABAergic neuron development - VIP | Mi et al. | | |
| **Zdhhc9** | CGE-Progenitor | Mi et al. | | |
| **Zeb2** | MGE-CIN | Chen et al. | | |
| **Zfhx3** | LGE-immature neurons | Chen et al. | Markers previously not found in GABAergic neurons | Chen et al. |
| **Zfhx4** | LGE-immature neurons | Chen et al. | | |
| **Zfp113** | GABAergic neuron development - PV | Mi et al. | | |
| **Zfp266** | MGE-immature neurons | Chen et al. | | |
| **Zfp367** | Progenitor | Mi et al. | | |
| **Zfp36l2** | GABAergic neuron development - SST | Mi et al. | | |
| **Zfp462** | MGE-immature neurons | Chen et al. | | |

| | | |
|---|---|---|
| **Zfp703** | dMGE-progenitor | Mi et al. |
| **Zfp715** | dMGE-progenitor | Mi et al. |
| **Zfp937** | dMGE-progenitor | Mi et al. |
| **Zfyve1** | GABAergic neuron development - PV | Mi et al. |
| **Zkscan4** | vMGE-CIN | Mi et al. |
| **Znrf1** | Progenitor SST-E14.5 | Mi et al. |

v: ventral; d: dorsal; PV: Parvalbumin; SST: Somatostatin; VIP: Vasoactive-intestinal peptide; CIN: Cortical Interneuron; CGE: Caudal ganglionic eminence; MGE: Medial ganglionic eminence; LGE: Lateral ganglionic eminence; E: embryonic day; Ref.: Reference source
